# A Paper-Droplet Hybrid System for Efficient Blood Plasma Separation in Multiplex Diagnostics

**DOI:** 10.1101/2025.01.14.632976

**Authors:** Amaan Dash, Rajeev Srikar Medicharla, Sunando DasGupta

## Abstract

Single μPAD-based lateral flow blood plasma separation techniques suffer from low sensitivity, inconsistent performance, and extended turnaround times, hindering point-of-care whole blood multiplexing. Integrating multifunctional microfluidic approach could offer a paradigm shift, providing rapid, high-throughput blood testing with reduced sample alongside streamlined downstream automation. Herein we report a novel low-cost sessile-drop and paper hybrid blood plasma separation technique that achieves efficient (≥ 98 %) separation within seconds (∼70 s). Evaporation-driven natural convection within a sessile blood droplet facilitates effective mixing, while capillary forces prevalent in the paper structure selectively transport plasma to the detection zones, all without any externally applied motive force. The separated plasma is successfully analysed for simultaneous glucose and albumin detection using colorimetric assay. This method demonstrates high accuracy in predicting glucose and albumin levels in healthy and diseased samples when benchmarked against an Automated Biochemistry Analyzer. A 3D foldable all-in-one paper microfluidic device, complemented by a MATLAB based application (GPS App.) has been developed for quantitative assessment during on-field diagnostics. This study presents a rapid, efficient, and user-friendly alternative to traditional flow-based separation, providing reliable detection in resource-limited settings.

## 1. Introduction

Paper platforms are extensively investigated as promising tools for point-of-care diagnostics, enabling timely and efficient detection across diverse applications. (Aghababaie et al., 2023), (Bezinge et al., 2024). It offers compact, cost-effective solutions for precise fluid manipulation facilitating rapid, accurate, and user-friendly testing in resource-constrained environments (Mukhopadhyay et al., 2022a), (Dash et al., 2024). The inherent self-assisted passive flow, along with disposability, and recyclability provides them competitive edge for on-field biomarker detection. Given these biomarkers are integrated with molecular signatures, there is burgeoning interest to perform multi-analyte testing for precise characterization of these specific disease states (Park et al., 2019) of an individual. However, limited spatial resolution, diffusion variability, cross-reactivity, and flow restrictions continue to restrict performance and compromise accuracy of multiplexed assays in paper analytical devices (μPADs). These challenges further exacerbate during blood-based analysis, which is routinely performed by clinicians to detect essential bio-analytes (Nilghaz and Shen, 2015), (Kim et al., 2020), (Al-Tamimi et al., 2022). Whole blood analysis has been particularly challenging because of intense colorimetric interference from erythrocytes, making preliminary plasma separation via gold standard centrifugation or sedimentation necessary. However, this demands expensive instrumentation, time and skilled personnel. Furthermore, pre-plasma separation for μPAD based testing undermines the wide range applicability of point-of-care devices. This underscores the necessity of the development of effective paper built in blood plasma separation unit to facilitate efficient on-field characterization of plasma biomarkers.

In pathology, plasma is examined to monitor blood glucose and protein levels among others, with glucose typically ranging between 70-100 mg/dL and protein 6-8 g/dL, with primarily albumin at 60 % (3.5-5 g/dL) (Pokhrel et al., 2020). Any deviation may indicate diabetes, kidney dysfunction, cardiovascular risk or underlying issues requiring further assessment. They are integral for disease management, and therefore clinicians frequently include them for general health screening. Hence, a μPAD platform for simultaneous glucose and protein detection from plasma could enable early identification of potential risks at low cost, even outside clinical settings. However, whole blood is primary analyte in outpatient scenario and therefore rapid and efficient plasma separation is instrumental for these testing. Prior techniques of utilizing patented membranes (Burgos-Flórez et al., 2022), (Guo et al., 2020), (Songjaroen et al., 2012), (Park et al., 2019), or agglutinating reagents in μPADs (Yang et al., 2012) can significantly inflate overall assay cost. RBC aggregation is a cost-effective yet efficient technique for separation by creating differential flow between cells and plasma within the paper matrix (Nilghaz and Shen, 2015), (Yang et al., 2012). However, aggregates formed can severely impede blood flow and increase turnaround time, reducing separation bandwidth (Kar et al., 2015), and compromising sensitivity of subsequent assay (Nilghaz and Shen, 2015), (Kim et al., 2020), (Al-Tamimi et al., 2022). Therefore, it is crucial to engineer novel techniques promoting readily integrable, more effective, and quicker plasma separation for μPAD based testing.

Utilizing sessile droplets for chemical and biological assays represents a transformative advancement, addressing inherent limitations of traditional flow-based systems (Garcia-Cordero and Fan, 2017), (Hernandez-Perez et al., 2016). Requiring only a hydrophobic substrate, they leverage internal flows within droplet during evaporation as efficient mixing strategy without external instrumentation (Barmi and Meinhart, 2014). Capillary and Marangoni flows are exploited for range of biomedical applications, including analyte concentration, particle separation, sorting, and cell-based assays (Yang et al., 2022). In blood droplet analysis, the focus is often exploiting after-effects of evaporation, such as drying patterns or crack formation (Parsa et al., 2018), which can be time-consuming. In contrast, directly leveraging hydrodynamics of evaporating blood droplet also holds significant potential to facilitate high-throughput biochemical and physiological reactions. Each sessile blood droplet function as small independent reactor with high surface-to-volume ratio and mixing, promoting uniform reaction condition. This can enable faster and highly efficient reaction kinetics involving hemoglobin, plasma components, and other critical constituents with low reagent/sample consumption to correlate with different medical conditions, whose implications are scarcely explored in literature. Consequently, its integration with μPAD platforms can combine precise microfluidic control, for rapid, low-cost, and efficient biomarker detection maintaining performance, stability, and accuracy across diverse array of biological assays.

In this work, we report a new sessile droplet-paper multifunctional microfluidic technique to achieve blood plasma separation (BPS) within few seconds, employing only commercially available grade-1 filter paper. We exploit evaporation as a naturally rapid and uniform mixing strategy and couple it with capillary forces inside the paper matrix for highly efficient plasma separation from blood, unmatched by conventional paper flow-based systems. The high-quality plasma obtained is analysed using MATLAB image processing techniques and subsequently utilized for detecting model targets i.e., glucose and albumin levels via colorimetric analysis. The paper device was calibrated using standard solutions; and the results for the whole blood and diseased samples were benchmarked against pathology-based automated reference technique for validation. The device was integrated into a 3D foldable paper platform, and a MATLAB desktop application has been developed for on-field analysis utilizing the proposed technique. The device could successfully predict glucose and albumin levels in healthy and diseased samples, serving as rapid and cost-effective solution for point-of-care hematological testing with quantitative assessment.

## 2. Materials and Methods

The blood samples were collected from individuals aged 20-60 years from the B. C. Roy Technology Hospital, Indian Institute of Technology Kharagpur, India. The samples were obtained through venepuncture into EDTA tubes, following standard laboratory protocol and after obtaining institutional ethical clearance (No. IIT/SRIC/DEAN/2022), maintaining confidentiality of participants and informed consent. All samples were used within 24 hr of collection. For details of the patient data; and methodology and materials used, please refer to Supplementary Material.

## 3. Results and Discussion

### 3.1. BPS Concept and Design Parameters

In the proposed technique, as illustrated in Fig. 1(a), red blood cell (RBC) aggregation is achieved within a non-movable sessile droplet, which is outside the paper matrix, followed by wicking of the solution mixture using a paper device. Firstly, a droplet of salt solution is dispensed to reside on a hydrophobic paper substrate (refer Fig. S1 for the substrate preparation) creating a base environment for aggregation kinetics to initiate, Fig. 1(a − I), followed by adding an equal volume of whole blood on top of the initial droplet. The density and surface tension difference between the two drops create strong internal currents, promoting rapid mixing of both the fluids. As the reactants are confined to a small volume (12 µL), the diffusion limitations are minimised and the time required for mixing and initiate aggregation is considerably reduced. After a time 𝑡 = 45 𝑠, the paper device was dipped vertically into the droplet consisting of blood and salt, as shown in Fig. 1(a − II). With the onset of cell aggregation, the pores of the paper device act as a barrier, restricting the movement of larger aggregates from the droplet into the paper device, while the plasma moves vertically up the device. The RBCs therefore get concentrated in and around the bottom of the paper strip and the separated plasma selectively wicks the paper to move towards the detection zone (discussed in more detail later in Fig. 2). The paper device was subsequently analysed under a microscope to quantify the plasma yield quality in the detection zone. This method effectively filters plasma from the whole blood with high quality and overcomes flow restrictions encountered in previous μPAD techniques, where drying agglutination/aggregation reagents on paper can reduce porosity, and aggregations induced within paper matrix can clog pores and block plasma movement. (Ardakani and Hemmateenejad, 2023)

**Fig. 1:**
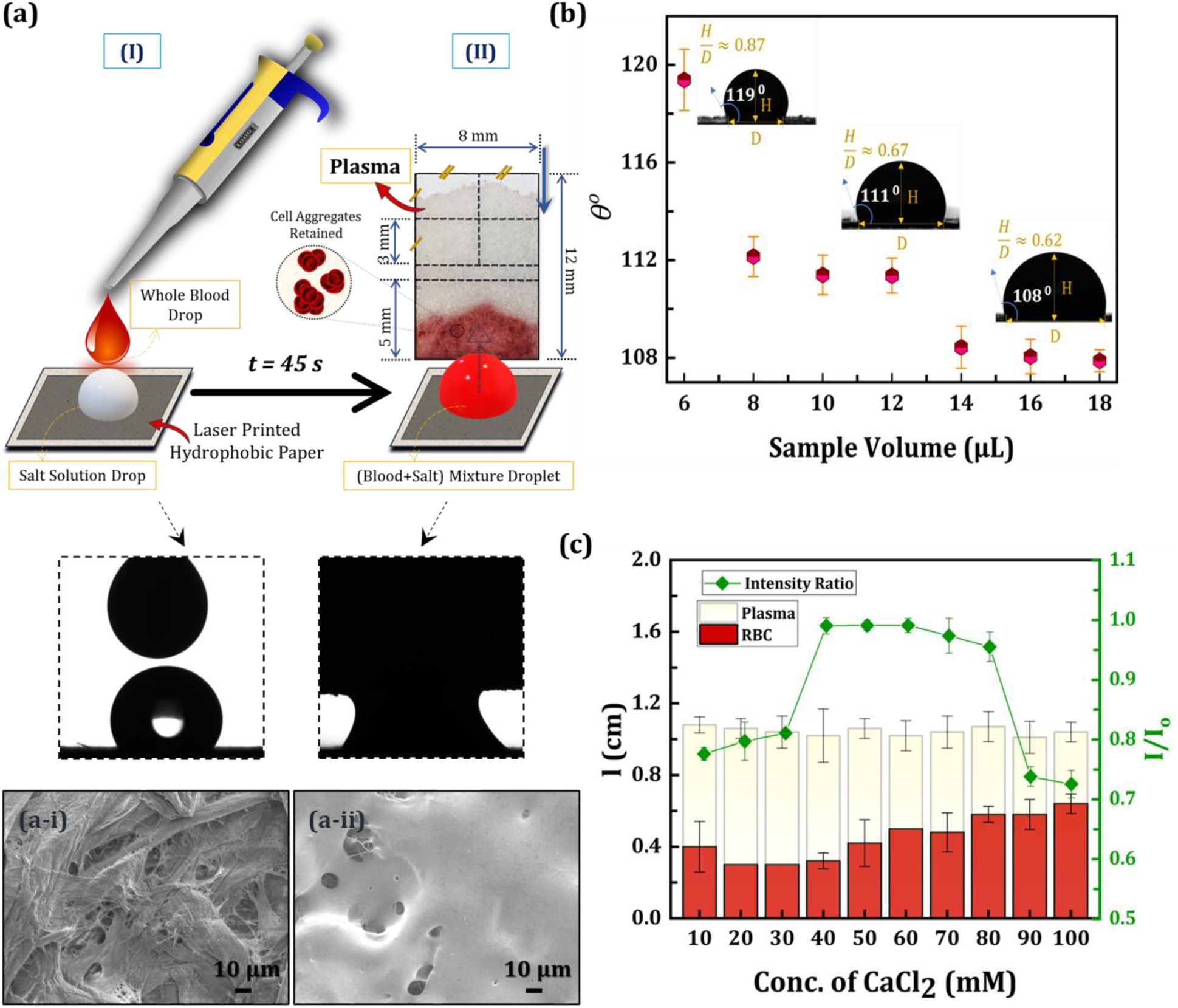
(a) Schematic of plasma separation procedure using sessile blood droplet’s internal convection and μPAD′s capillary transport, where 𝑡 = 45𝑠 is the time for effective cell aggregation. The SEM images of: (a-i) un-modified grade-1 filter paper, and (a- ii) laser printed paper substate, provide clearer difference of the changes. **(b)** Variation of aspect ratio 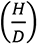 and contact angle 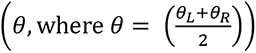 of the droplet with varied sample volumes (in 1:1 ratio salt and blood) (mean ± SD, 𝑛 = 5). The contact angles were obtained using a Goniometer, (Data Physics, OCA 15 Pro), and aspect ratios using ImageJ. **(c)** Distance, 𝑙(mm) traversed by RBCs & separated plasma after complete drying, 𝑡 ≈ 10 min; and separation extent 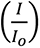 at 𝐼_𝑜_ detection zone for varied concentration of CaCl_2_ (mean ± SD, 𝑛 = 5).

**Fig. 2.**
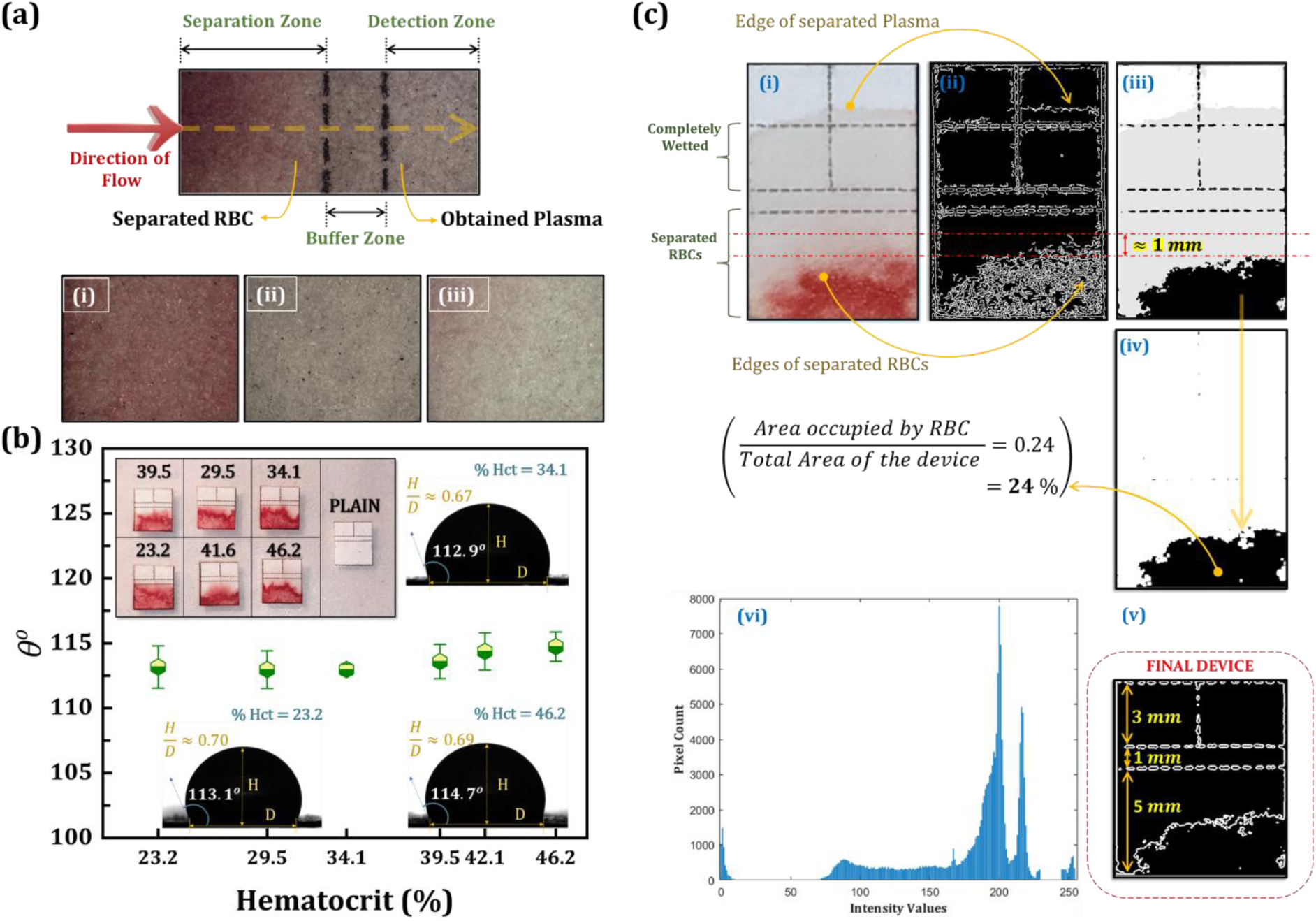
(a) Microscopic images (Leica DM6000M) of μPAD zones after plasma separation using 5× objective lens. The images of the detection zone for: (i) Blood with added 1:1 DI water (completely hemolysed blood) (ii) Blank device without any sample, and (iii) Added 1:1, 40mM CaCl_2_ mix blood sample. **(b)** Contact angle (𝜃^𝑜^) of sample droplet for varied blood hematocrit (Hct %) showing marginal change. The inset image highlights actual paper strips for different blood hematocrit concentrations. **(c)** MATLAB image processing protocol: (i) Image of actual BPS paper device processed, (ii) Canny’s Edge Detection, (iii) Otsu’s Thresholding, (iv) Fractional total area coverage of RBCs, (v) Perimeter Detection of optimised design, and (vi) Histogram of intensity values for actual device image after converting to grey scale.

The separation is further augmented by utilizing grade-1 filter paper as dipstick (average pore size ∼11 μm), considering blood cell dimension range from ∼6−14 μm, to effectively filter cell aggregates. The dipstick dimensions were optimized to maximize the separation bandwidth, (Δ𝑙_𝑠𝑒𝑝._ = 𝑙_𝑝𝑙𝑎𝑠𝑚𝑎_ − 𝑙_𝑅𝐵𝐶_) (refer Table S1 and Fig. S2). The pore size in the range of blood cell facilitates larger Δ𝑙_𝑠𝑒𝑝._ and maintain higher flowrate for faster separation. Grade-1 is also relatively low-cost, lack flow rate-hindering additives, and has uniform pore distribution (Giokas et al., 2014), promoting consistent colorimetric assay (Ardakani and Hemmateenejad, 2023). Paper is chosen here as the model substate for droplet placement due to its easily modifiable physical/chemical properties (Modaressi and Garnier, 2002) allowing integration with pre-existing μPAD designs. Since, liquid absorption inhibit droplet formation on paper, a grade - 1 filter paper, Fig. 1(a-i), was patterned hydrophobic, as shown in Fig. 1(a-ii), to promote hemophobicity (Dash et al., 2024). To preliminarily validate blood plasma separation and assess repeatability, tests were conducted intra-day with 50 different blood samples (Fig. S3−S5, Table S2 and Supplementary Video S1). Detection zones were excised and tested by immersing into standard Bromocresol Green (BCG) laboratory reagent, producing a colour change from yellow to blue, thereby confirming presence of plasma in detection zones (refer Fig. S6).

The droplet shape plays a critical role in achieving rapid and efficient separation. The shape blood droplet assumes on paper results from the combined influence of surface- fluid interaction, paper roughness, and gravity. As shown in Fig. 1(b), with increase in sample volume, the contact angle of the sessile blood droplet decreased. This decrease follows a stick-and-jump transition between different sample volumes due to contact line pinning into roughness of paper (Modaressi and Garnier, 2002). The aspect ratio i.e. 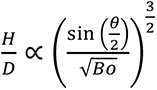 of the sample droplet similarly decreased because of decrease in contact angle (𝜃) and increase in Bond Number 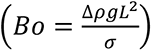 (Lubarda and Talke, 2011). This leads to sample droplet transition from spherical cap to oblate spheroidal0 shape (ellipsoidal), a trait undesirable in this context as it diminishes sample repellence of the patterned paper. Higher sample repellence will allow plasma to easily mitigate towards the top, thereby promoting higher capillary suction into paper strips. Also, maintaining a higher contact angle results in higher Marangoni number, which increases the strength of recirculating currents and enables better convective mixing for faster aggregation inside blood droplet mixture (Barmi and Meinhart, 2014), (Trantum et al., 2014). Additionally, a spherical cap minimizes surface area, preventing evaporative losses and resulting in greater extent of separated plasma movement in paper channels. Therefore, a 12 μL sample droplet (6 μL (blood) + 6 μL (salt solution)) ensured sufficient plasma separation while simultaneously retaining a spherical shape, making it optimal for further analysis.

### 3.2. Effect of Salts on BPS

Investigating the effect of salts is instrumental for maximising red blood cell (RBCs) aggregation within the droplet while concurrently regulating hemolysis. The reddish appearance of the separated plasma, stemming either from improper separation or release of haemoglobin can severely interfere with precise colorimetric analysis. Hence, to maximize plasma separation without hemolysis in minimal time, four distinct salts: NaCl, KCl, CaCl2 and MgCl2 were examined across varied concentrations (10 − 100 mM), see Fig. S7(a). Notably, salts containing higher metal ion valencies demonstrated higher separation extent 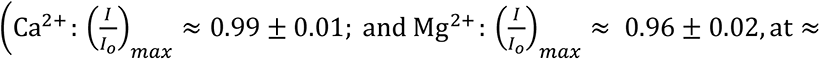 40 − 60 mM). A higher counter valency (𝑧) increases the charge density around RBCs consequently supressing electric double-layer repulsion (Kontogeorgis and Kiil, 2016). This is also evident from the Hardy-Schulze rule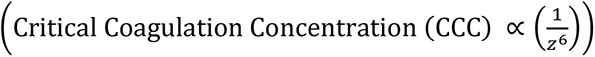 (Nilghaz and Shen, 2015).

Furthermore, calculating the Debye Length 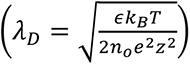 by considering approximate dielectric constant (𝜖) ≈ 60𝜖_𝑜_ for blood droplet (Gabriel et al., 1996) at room temperature (𝑇) ≈ 37℃ with a 40 − 60 mM salt concentration, we obtain 𝜆_𝐷,1+_(≈ 389 ∓ 32 nm) > 𝜆_𝐷,2+_(≈ 194 ∓ 16 nm). This confirms the significance of counter-ion valency in reducing electrostatic interaction of RBC suspension inside the droplet mixture. Following the same trend, a salt with trivalent ion, 𝜆_𝐷,3+_(≈ 129 ∓ 11 nm) could be utilised. However, CaCl2 (𝑧 = +2) is utilised in this study because Ca^2+^ is present as a natural and essential component of physiological blood coagulation process in human body that ensures proper functioning of vital blood processes. It also serves as an excellent exogeneous coagulation factor capable of indirectly activating platelets (Toyoda et al., 2018). This in conjunction with charge shielding is expected to promote formation of thick, well cross-linked fibrins inside the blood droplet mixture, further adding to better RBC aggregation in vitro.

For varied concentrations of CaCl_2_solution, the extent and bandwidth of separation is shown in Fig. 1(c). The percentage plasma yield quality (𝜂) in the detection zones of the paper device was computed to estimate the levels of hemolysis and effectiveness of separation. Due to hypotonic effects, the mean grey intensity ratio of paper devices obtained from blood droplet mixed with deionized (DI) water, instead of salt solution, and subjected to wicking is considered as the measure of 100% blood hemolysis in the detection zone (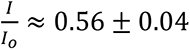; see Fig. 2(a − i) and Fig. S7(b)). This value is used as 𝐼_𝑜_ control, (𝐼𝑅_𝑐𝑜𝑛𝑡𝑟𝑜𝑙_) and all intensity ratios are considered relative to the blank paper device (𝐼_𝑜_) ⇒ 𝐼𝑅_𝑏𝑙𝑎𝑛𝑘_ = 1, Fig. 2(a − ii). Hence, percentage yield quality is computed as a fraction of control using the following modified expression:

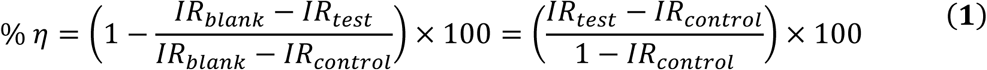

The calculations presented in Table S3 suggests that quality of plasma (𝜂) obtained along with maximum separation bandwidth (Δ𝑙_𝑠𝑒𝑝._) peaks at an intermediate salt concentration of 40 mM, while reddish shades are noticeable in detection zones at both higher and lower levels. At low salt concentrations (0 − 30 mM), a hypo-osmotic environment outside cell membrane may be established (Nepal and Rao, 2011). The hypo-osmotic stresses could induce morphological alterations in red blood cells (RBCs). This could cause an increased influx of Ca^2+^ions and solution into the cells, resulting in hemolysis (𝜂 ≈ 48 − 56%). Similarly, at higher concentrations of salt (≈ > 80 mM), the hyper-osmotic environment causes RBC to shrivel. This might result in a greater RBC transport in paper channel reducing bandwidth (Δ𝑙_𝑠𝑒𝑝._ ≈ 0.4 cm) and plasma quality (𝜂 ≈ 36 − 39 %) at the detection zones. At recorded 40 mM CaCl_2_, an iso-osmotic condition across RBCs likely yielded the most favourable separation, with plasma quality of 𝜂 ≈ 97.7 ± 3 % and separation bandwidth, Δ𝑙_𝑠𝑒𝑝._ ≈ 0.7 cm, Fig. 1(c) and Fig. 2(a − iii), wherein separated plasma exhibited distinct slightly yellowish tint, representing true colour of blood plasma.

The plasma yield quality is heavily reliant on the separation time. Faster separation not only improves turnaround time for clinical tests but also help circumvent issues of clogging, evaporation, and influx of cells into the detection zone. The droplet-dipstick technique herein offers flexibility to promptly remove the paper device from over the sessile droplet, preventing cell leakage and achieving high quality plasma with higher separation bandwidth. The total separation time (𝑡_𝑠_) is analysed as cell aggregation time (𝑡_𝑎_) and paper strip wicking time (𝑡_𝑤_): 𝑡_𝑠_ = 𝑡_𝑎_ + 𝑡_𝑤_. Rapid mixing in the small droplet volume facilitates faster aggregation, with separation peaking at 𝑡_𝑎_ ≈ 45 𝑠, (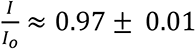, see Fig. S8(a)). To ensure complete wetting of detection zones by plasma while preventing any cell leakage and backflow, the paper strip contact time with the droplet was limited to 𝑡_𝑤_ ≈ 25 𝑠, see Fig. S8(b). It effectively eliminated the risk of cross-contamination of reacting species and development of unintended colours, alongside preventing influx of RBC into the detection zone. This resulted in a total blood/ plasma separation time of 𝑡_𝑠_ = 70 𝑠 (see Fig. S8(c)) which is significantly faster compared to ≈ 5 − 10 min required by gold-standard laboratory centrifuge. The assay time is appreciably less, negating the loss due to evaporation. Consequently, the proposed technique can be performed under varied on-field environmental conditions without the need for temperature or humidity control.

### 3.3. Plasma Separation Efficiency

The blood plasma separation efficiency is quantified as the ratio of separated plasma volume recovered to the entirety of plasma introduced into the device. Firstly, blood hematocrit levels in the range of 30-50% (average Hct ≈ 40±10%) were considered for the analysis. This corresponds to ≈ 60 ± 10 % plasma fraction, equivalent to ≈ 3.6 ± 0.6 μL of plasma for optimised 6 μL blood sample dispensed. With addition of 40mM CaCl_2_solution in 1:1 ratio, ≈ 9.6 ± 0.6 μL of diluted plasma is present in a total 12 μL sample mixture examined. Secondly, the separated plasma in the paper device was quantified using the method described by (Ardakani and Hemmateenejad, 2023). The recovered plasma volume was calculated to be ≈ ≥ 9.46 ± 0.52 μL in the paper device. Thus, the proposed technique demonstrates a high plasma separation efficiency of ≥ 98.5 ± 0.1 %, which can be seen in Fig. 1(a) and Table S2, and remained consistent throughout the entire study.

### 3.4. Hematocrit Influence on BPS

One critical aspect in point-of-care blood analysis is maintaining consistently high separation efficiency and yield quality across broad blood types. Blood hematocrit (Hct) could severely impact and delay flow-based blood plasma separation techniques (Songjaroen et al., 2012); therefore, its influence on plasma separation has also been evaluated. The aspect ratio 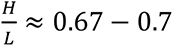, and contact angle (𝜃 = 𝜃 = 𝜃 ≈ 112 − 114^𝑜^) of the droplets did not show notable differences for varied haematocrit levels, see Fig. 2 (b) and Fig. S4 . Firstly, although an increase in hematocrit is expected to reduce the contact angle of sessile blood droplet (Hct ↑ ⇒ 𝜃_𝑐_ ↓) (Pitts et al., 2013), a hydrophobic porous substrate is strategically implemented here to induce metastable pinned state across blood sample with varying RBC fractions. The reduction in interfacial energy was insufficient to overcome pinning effect of sessile droplet, causing droplet shape to remain mostly unaffected by blood hematocrit. Secondly, the capillary driving force of dipstick generates higher initial transport rate for the liquid component of the blood mixture. It is minimally influenced by the RBC fraction, as they remain outside the paper matrix and experience delayed transport due to drag from liquid uptake by the paper. Therefore, the plasma flow and separation restriction that could occur due to clogging, if aggregation were induced within the paper matrix, especially at higher hematocrit, are largely minimized in this technique. Consequently, due to the stable pinned state and pre-induced aggregation inside the droplet, consistently high quality and efficiency of separation is observed irrespective of blood haematocrit.

### 3.5. Simultaneous Glucose and Protein Detection

The developed droplet and paper dipstick technique is utilised for direct determination of glucose and albumin levels in the whole blood. Successful biomarker detection of interest from plasma is not only a clear indication of effective separation but also establishes applicability of the technique as a point-of-care testing platform. However, due to the diffuse or tapered leading front of RBC during blood flow in the paper matrix, manually determining the exact length traversed by the cells could be prone to human errors, as shown in Fig. 2 (c − i). Therefore, MATLAB image processing techniques were implemented to accurately quantify the traversal length, free of any personal bias. The detection zones for analyte detection were then designed and marked on the paper device, ensuring precise and reliable measurements. The maximum length (representing both the core and diffuse RBC front) and minimum length (representing only the core RBC front) were obtained using MATLAB edge detection, Fig. 2 (c − ii), and thresholding techniques, Fig. 2 (c − iii). An approximate 1 mm difference constitutes a diffuse RBC layer, highlighting the sharpness of transition, which may vary with blood hematocrit, flow dynamics and paper properties. This is quite difficult to control and a 1 mm extra length is therefore added as an additional safety margin to the characterization process. This approach is crucial because while the darker RBC shades are prominent initially, the diffuse layer could interfere with colorimetric assay as the device dries.

The darker RBC front, with higher cell concentration, covers a minimum of ≈ 24 − 27 % of total area (minimum is considered to set practical and resource efficient safety limits), Fig. 2 (c − iv) and Table S4. Therefore, the separation zone was kept 1.5 times this area (a factor of 1.2 − 1.5 is a common engineering safety margin). This corresponds to ≈ 40 % of total area i.e., ≈ 5 mm of the total 12 mm length. To account for the diffuse RBC layer (as discussed previously), an additional 1 mm length was added, making the total separation zone approximately 6 mm. The separated plasma could completely wick an additional 3 mm length, Fig. 2 (c − v)), following this zone within a time span of 70 s. Therefore, it is prudent to choose this design, after removing any excess sections ensuring that the zone was equally segmented for multiplexing. The highest pixel count is also concentrated around high intensity values, Fig. 2 (c − vi) indicating that separated RBCs are confined to a small fraction of the total channel and the plasma obtained is of high quality. Specific reagents (refer Supplementary Material S6) are dried in the respective detection zones and the concentration of glucose and albumin were obtained using colorimetric assay. The modified paper strip was immersed into the final droplet consisting of 6 μL (40 mM CaCl_2_solution) + 6 μL (standard solution of dextrose or human serum albumin (HSA); or whole blood) for BPS followed by simultaneous detection. The schematic of the assay procedure is shown in Fig. 3 (a).

**Fig. 3.**
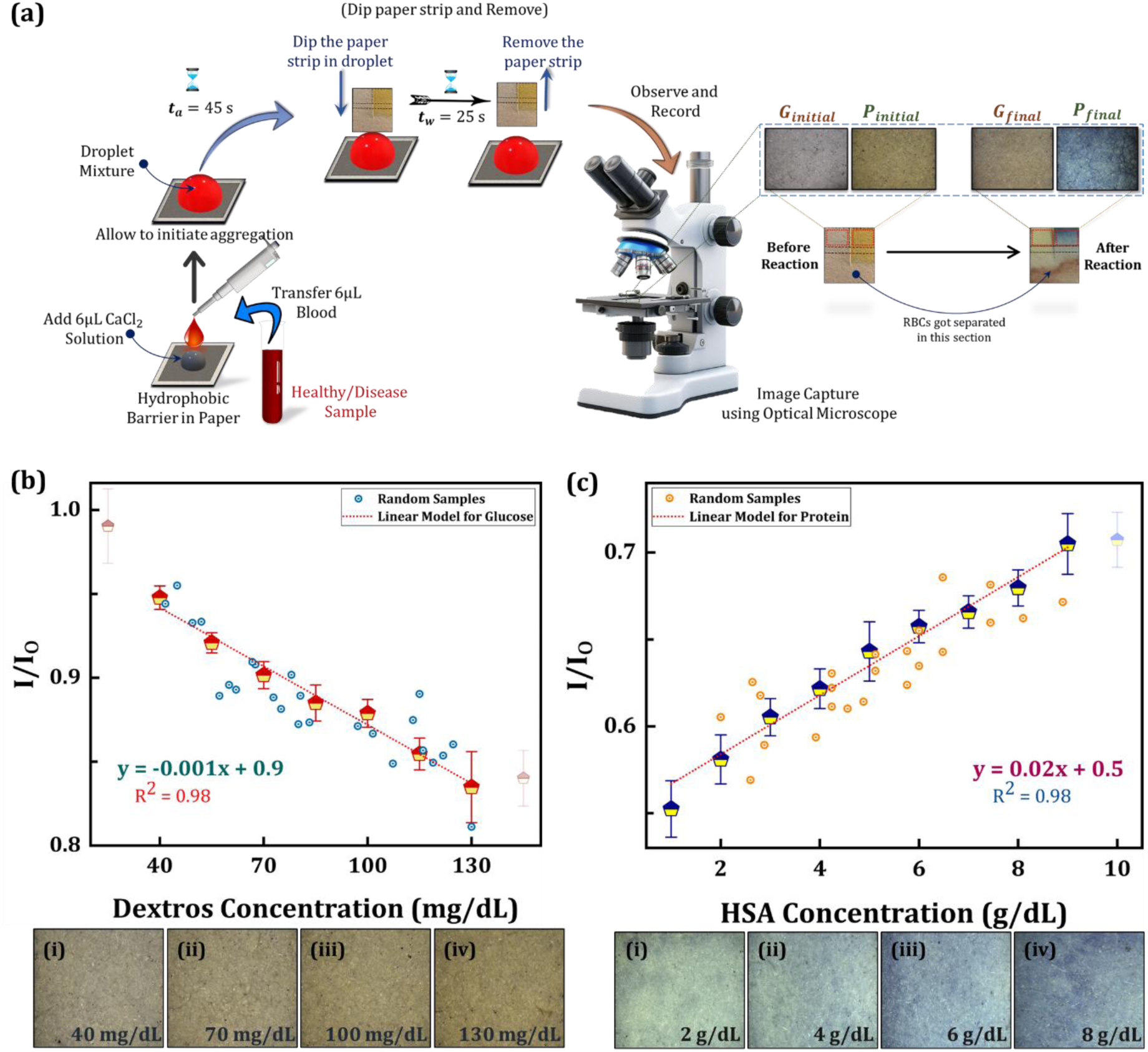
(a) Schematic representation of blood plasma separation process and subsequent analysis from standard solutions/healthy/diseased samples. Standardization was performed using images captured by Optical Microscope (Leica DM6000M) and analysed using ImageJ. The actual paper device design is highlighted, featuring a central cut to divide fully wetted region into two equal halves, enabling simultaneous glucose and albumin detection through colorimetric assays. **(b)** & **(c)** Linear calibration curve correlating colour intensity ratio 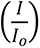 vs. concentration of 𝐼_𝑜_ analyte in standard dextrose and HSA solutions respectively (mean ± SD, 𝑛 = 15). Statistical analysis, including ANOVA and t-tests was performed for all data. Additional testing was conducted with 𝑛 = 25 random concentration (circular scatter points) within the linear range, and the results were validated using Student’s t-test.

#### 3.5.1. Device calibration for muti-analyte assay

To standardize the detection method, glucose and albumin concentration in standard dextrose and human serum albumin (HSA) samples were detected and results were used to obtain the calibration curves for each. A well-defined linear calibration curve correlating colour intensity ratio 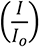vs. concentration of analyte is observed, for glucose ranging from 40-130 mg/dL and for albumin ranging from 1-9 g/dL. The paper sensor colour changed from colourless to golden brown for glucose, and yellow to blue for albumin, in their respective detection zones, see Fig. 3 (a). The mean grey intensity value for glucose and blue channel intensity value for albumin provided the best results and are therefore used for evaluating intensity of the colour developed. A linear regression model with 95% confidence level is fit to experimental data for glucose with 𝑅^2^ = 0.983, Fig. 3 (b) and protein with 𝑅^2^ = 0.980, Fig. 3 (c). The LOD (blank + 3𝜎) is 5.8 mg/dL and 0.7 g/dL; and COV is 2.4 % and 6.6 % respectively for glucose and albumin which shows sufficient detection ability for analyte detection. ANOVA performed on both linear models resulted in p-values = 1.78 × 10^−5^ (for glucose) and 2.89 × 10^−7^ (for albumin), both of which are < (𝛼 = 0.05). This concludes that linear regression model is significant for both the analytes. The t-statistics value, with < (𝛼 = 0.05), further confirms that the slope is significant, indicating colour intensity ratio is a significant variable to differentiate between the two concentration ranges, refer Fig. S9. To further validate the curve and ensure precision in future estimations, additional 25 random concentrations from the linear range were tested, conducting an individuality test. The COV is ≈ 1.82 % for glucose, Fig. 3 (b); and ≈ 2.50 % for albumin, Fig. 3 (c); scatter points. It is assumed that randomly selected concentrations follow t-distribution. A Student T-test is performed to compare experimentally obtained intensity ratio (IR) with that predicted from regression model. For two-tail test, the obtained p-values = 0.88 and 0.54, for glucose and albumin respectively, both > (𝛼 = 0.05). Therefore, null hypothesis (𝐻_𝑜_) i.e. random data points follow their corresponding calibration curve can be accepted, see Fig. S10.

#### 3.5.2. Simultaneous Glucose and Albumin detection

For initial validation of the accuracy and test multiplexing capabilities of the proposed method, the glucose and albumin levels were measured simultaneously in a single device using 5 healthy whole blood samples (refer Table S5-S6 and Fig. S11). Each sample was tested in triplicate for glucose and albumin level detection, with relative standard deviation (RSD) ≤ 5 %. The detection accuracy was evaluated by comparing results with training data from standard solutions. A 95 % confidence and 95 % prediction band for calibrated glucose and albumin linear regression model were obtained using Eq. (2) and Eq. (3), as follows:

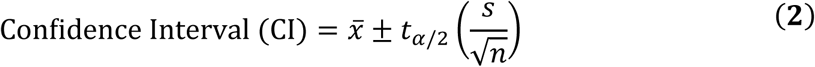

and

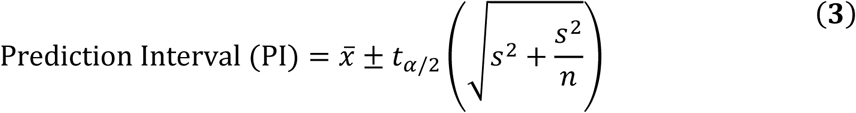

where 𝑥̅ is mean intensity obtained, 𝑡_𝛼/2_is obtained from t-distribution for 95 % based on degrees of freedom (𝑑𝑓 = (𝑛 − 1)), 𝑠 is sample standard deviation and 𝑛 is sample size. The device could successfully predict glucose and protein levels within 95 % prediction interval when validated against an Automated Biochemistry Analyzer (DiaSYS 400).

To conduct a more comprehensive analysis of the detection capabilities, the accuracy was further validated with additional 30 healthy whole blood, 30 diluted and 30 spiked samples (refer Table S7) to cover broader sample spectrum from which glucose and albumin levels were detected simultaneously. The samples were spiked by fixed 1.2 g/dL standard HSA solution. The device could successfully predict glucose and albumin levels as shown in Fig. 3 (a) and Fig. 3 (b) with ≈ 85 % and ≈ 89 % sensitivity respectively. This underscores precision and versatility in accurate multiplexing across a broad sample spectrum. Furthermore, to evaluate the potential cross-interference between glucose and albumin assay, we calculated Pearson’s correlation coefficient between both the intensity ratios for all blood samples. A coefficient of ≈ 0.054 (𝑅^2^ = 0.003), (see Fig. S12) indicates no cross-dependence between the assays. Although no straightforward relationship was observed, small dependence could be patient-specific. Therefore, these associations should be interpreted within broader clinical and metabolic context of individual patient.

#### 3.5.3. Testing with diseased samples

To further assess the accuracy of testing method, the device was evaluated using whole blood samples from patients with varying glycemic and albuminemic conditions, including hyper (100 − 125 mg/dL)/ hypo (54 − 70 mg/dL) glycaemia, hyper (5 − 6.5 g/dL) / hypo (≤ 3.5 g/dL) albuminemia, and Diabetes Mellitus. Mild glycaemia and albuminemia do not impart immediate complications and can often be managed through lifestyle modifications. Extreme levels or diabetic conditions necessitate medical intervention and regular follow-up, emphasizing critical need for rapid, reliable, and low-cost point-of-care estimation methods. The proposed plasma separation technique effectively detected glucose and albumin levels in diseased samples, achieving successful differentiation of specific conditions with ≥ 90 % sensitivity. Validation against the Automated Biochemistry Analyzer (DiaSYS 400), as summarized in Table 1, confirms the robustness of the method. Thus, the proposed approach serves as a promising platform for preliminary disease screening, enabling early health interventions.

**Table 1.** Plasma separation and simultaneous glucose and albumin level detection were performed for 15 diseased patients. Results obtained using current method were compared with reference values reported from DiaSYS 400 automated system.

| SR. No. | Hematocrit (%) | Glucose Reported (mg/dL) | Glucose Predicted (mg/dL) | Total Protein (g/dL) | Albumin Reported (g/dL) | Albumin Predicted (g/dL) | Current Method Range Obtained |  | Condition Reported |
| --- | --- | --- | --- | --- | --- | --- | --- | --- | --- |
|  |  |  |  |  |  |  | Glucose | Albumin |  |
| 1. | 41.1 | 107 | 100-130 | 7.9 | 4.9 | 3.5-5 | High | Normal | Hyperglycaemia |
| 2. | 34.1 | 113 | 100-130 | 5.9 | 3.5 | 3.5-5 | High | Normal | Hyperglycaemia |
| 3. | 39.4 | 238 | > 130 | 6.6 | 4.1 | 3.5-5 | Very High | Normal | Diabetes Mellitus |
| 4. | 38.8 | 118 | 100-130 | 5.9 | 3.6 | $\leq 3.5$ | High | Low | Hyperglycaemia |
| 5. | 38 | 346 | > 130 | 7.6 | 4.3 | 3.5-5 | Very High | Normal | Diabetes Mellitus |
| 6. | 34.8 | 140 | > 130 | 7.7 | 4.4 | 3.5-5 | Very High | Normal | Diabetes Mellitus |
| 7. | 25.9 | 101 | 70-100 | 5.8 | 3.2 | $\leq 3.5$ | Normal | Low | Hypoalbuminemia |
| 8. | 38.6 | 291 | > 130 | 6.1 | 4.4 | 3.5-5 | Very High | Normal | Diabetes Mellitus |
| 9. | 29.4 | 127 | 100-130 | 5.9 | 2.4 | $\leq 3.5$ | High | Low | Hypoalbuminemia & Hyperglycaemia |
| 10. | 30.9 | 91 | 100-130 | 5.8 | 3.4 | 3.5-5 | High | Normal | Hypoalbuminemia |
| 11. | 33 | 114 | 100-130 | 6.6 | 3.9 | 3.5-5 | High | Normal | Hyperglycaemia |
| 12. | 37.6 | 159 | > 130 | 6.6 | 4.1 | 3.5-5 | Very High | Normal | Diabetes Mellitus |
| 13. | 39.9 | 137 | > 130 | 7.2 | 4.2 | 3.5-5 | High | Normal | Hyperglycaemia |
| 14. | 37.6 | 178 | > 130 | 6.7 | 4.1 | 3.5-5 | Very High | Normal | Diabetes Mellitus |
| 15. | 45.8 | 125 | 100-130 | 7 | 4.6 | 3.5-5 | High | Normal | Hyperglycaemia |

### 3.6. Integration and Automation

#### 3.6.1. Device Integration

To improve user-friendliness and point-of-care applicability, the hydrophobic region and test strip were integrated into a single, compact, and foldable paper microfluidic platform. This device was fabricated using a low-cost tabletop laser printer, facilitating easy fabrication for mass production (Mukhopadhyay et al., 2022b). The base was reinforced with a 3 mm thick double-sided tape to improve mechanical durability, making the device more resistant to bending and twisting during handling. The 3D design accommodates droplet placement, plasma separation, and analyte detection (glucose and albumin) within a unified platform, as illustrated in Fig. 5(a). Its flexible, foldable design facilitates convenient transportation and storage, while the integration offers a scalable and practical solution for biomarker detection using the proposed separation method.

**Fig. 4.**
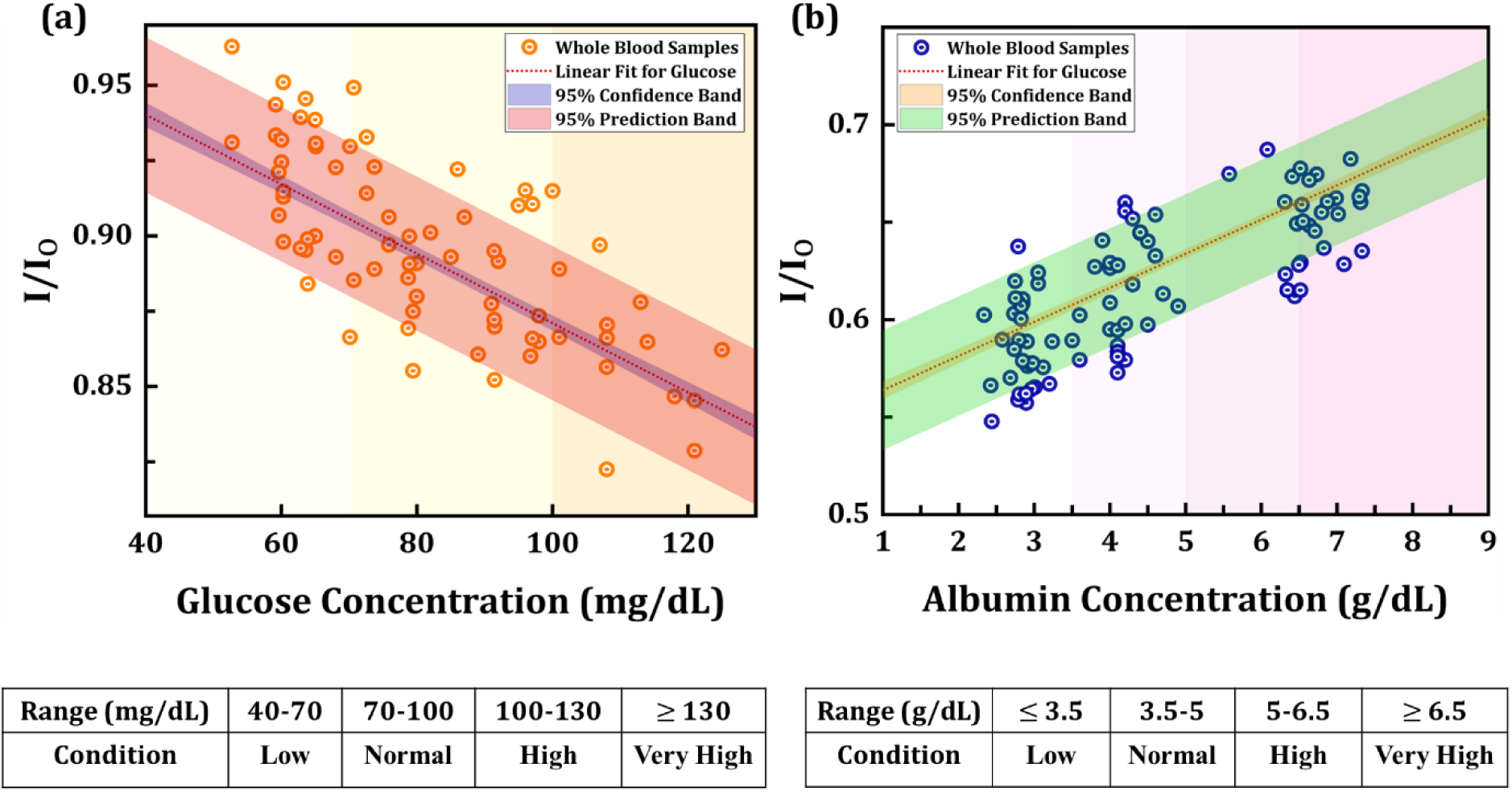
(a) & **(b)** Results of glucose and albumin detection post separation obtained from 𝑛 = 90 samples (= 90 × 2, each for glucose and albumin, totalling 180 tests; including whole blood, diluted and spiked samples). The linear regression model developed from calibration data is also plotted alongside 95% confidence and prediction bands. For practical on-field assessment, the concentration range was stratified into defined intervals. Further details regarding the blood samples are provided in Supplementary Material S9.

**Fig. 5:**
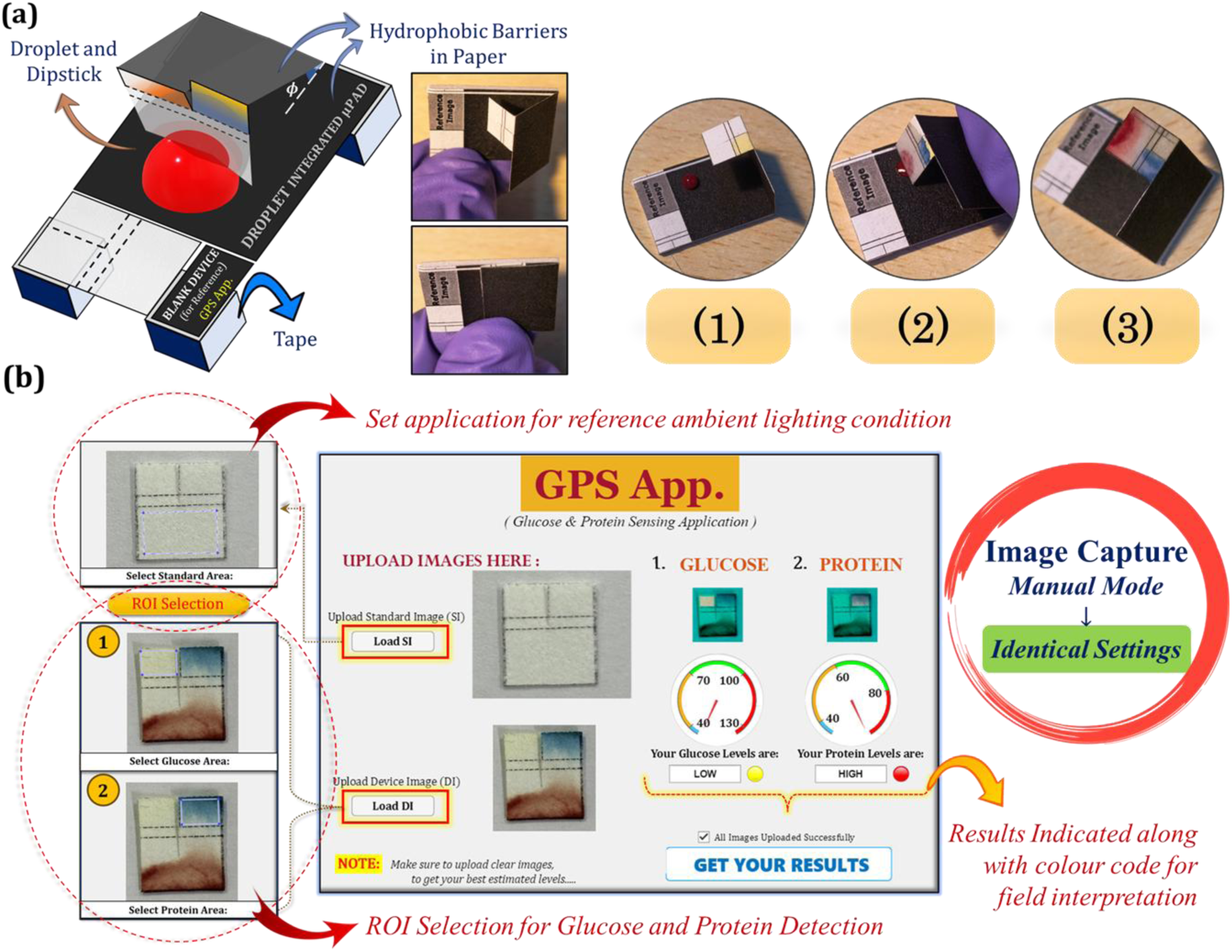
(a) A 3D foldable and integrated paper device designed with dedicated blank, hydrophobic, and detection zones for enhancing scalability and user-friendliness. This configuration provides stability and facilitates automation, making it ideal for point-of-care testing. **(b)** Graphical User Interface (GUI) of developed MATLAB (R2017b) Desktop Application (GPS App.), with region of interest (ROI) selection for multiplexing in whole blood samples, streamlining the testing process.

#### 3.6.2. In-house Application

To elevate practical usability of proposed technique for simultaneous separation and multiplexing, an automated Glucose and Protein Sensing (GPS) Application has been developed using MATLAB (R2017b). Herein, user initially loads a standard image (SI) of blank device to set application for reference ambient lighting condition. Subsequently, user uploads device image (DI). Once both images are uploaded, the application will prompt to select region of interests (ROIs) for blank, glucose and protein, see Fig. 5(b). Upon successful selection, respective ROIs will be delineated in glucose and protein sections respectively and application calculates individual intensity ratios. These ratios are then mapped to concentration ranges based on pre-loaded, experimentally established calibration data. The application estimates extent of colour grade and categorizes biomarker level, which is displayed to user through an edit-field output and lamp indicator. The robustness under varying lighting conditions is demonstrated in Supplementary Video S2, which confirms consistent performance in predicting concentration intervals. This combined with successful blood plasma separation and multiplexing, provides a reliable, quantitative tool for on-field diagnostics, thereby advancing point-of-care testing capabilities (refer Supplementary Material S10).

## 4. Conclusion

This study develops a first of its kind technique leveraging hydrodynamics of sessile droplets for rapid and efficient paper-based blood plasma separation, positioning it among the fastest existing paper or thread-based techniques. By synergizing droplet natural convection currents with capillary forces of paper, this technique achieves ≥ 98 % plasma separation efficiency with ≥ 95 % plasma yield quality from small samples (∼6 μL) in just ∼70 s, independent of the hematocrit levels. Multiplexing evaluations demonstrated ≈ 85 % and ≈ 89 % sensitivity in predicting glucose and albumin levels in whole blood, and spiked samples, along with ≥ 90 % sensitivity in diagnosing specific diseased states when validated against an Automated Biochemistry Analyzer. A 3D foldable paper-microfluidic platform, integrated with a custom image processing application, enables on-field detection with minimal human intervention. We envision this method to combine efficiency, precision, and versatility for rapid, low-cost diagnostics, thereby bridging the gap between research and point-of-care applications.

## CRediT authorship contribution statement

**Amaan Dash**: Conceptualization, Data curation, Formal analysis, Visualization, Methodology, Software, Writing – Original draft; **Rajeev Srikar**: Data curation, Methodology, Formal analysis; **Sunando DasGupta**: Investigation, Project administration, Funding acquisition, Supervision, Writing – review & editing

## Acknowledgements

The authors gratefully acknowledge the financial support provided by the Indian Institute of Technology Kharagpur, India [Sanction Letter no.: IIT/SRIC/ATDC/CEM/2013-14/118, dated 19.12.2013]. AD is thankful to the Ministry of Education, Government of India and IIT Kharagpur for the Prime Minister’s Research Fellowship (PMRF). We acknowledge the co-operation of Pathology division of the B. C. Roy Technology Hospital, Indian Institute of Technology Kharagpur, India for providing blood samples in accordance with Institute’s Ethical Guidelines as well as for providing gold standard biochemistry glucose and albumin analysis data for healthy and diseased samples.

## References

1. Aghababaie, M., Foroushani, E.S., Changani, Z., Gunani, Z., Mobarakeh, M.S., Hadady, H., Khedri, M., Maleki, R., Asadnia, M., Razmjou, A., 2023. Recent Advances In the development of enzymatic paper-based microfluidic biosensors. Biosens. Bioelectron. 226, 115131. 10.1016/j.bios.2023.115131

2. Al-Tamimi, M., Altarawneh, S., Alsallaq, M., Ayoub, M., 2022. Efficient and Simple Paper-Based Assay for Plasma Separation Using Universal Anti-H Agglutinating Antibody. ACS Omega 7, 40109–40115. 10.1021/acsomega.2c04908

3. Ardakani, F., Hemmateenejad, B., 2023. Pronounced effect of lamination on plasma separation from whole blood by microfluidic paper-based analytical devices. Anal. Chim. Acta 1279, 341767. 10.1016/j.aca.2023.341767

4. Barmi, M.R., Meinhart, C.D., 2014. Convective flows in evaporating sessile droplets. J. Phys. Chem. B 118, 2414–2421. 10.1021/jp408241f

5. Bezinge, L., Shih, C.J., Richards, D.A., deMello, A.J., 2024. Electrochemical Paper-Based Microfluidics: Harnessing Capillary Flow for Advanced Diagnostics. Small 2401148, 1–20. 10.1002/smll.202401148

6. Burgos-Flórez, F., Rodríguez, A., Cervera, E., De Ávila, M., Sanjuán, M., Villalba, P.J., 2022. Microfluidic Paper-Based Blood Plasma Separation Device as a Potential Tool for Timely Detection of Protein Biomarkers. Micromachines 13. 10.3390/mi13050706

7. Dash, A., Mukhopadhyay, M., Shaw, J., Bhattacharya, M., DasGupta, S., 2024. Rapid hematocrit estimation using a fold-crease induced fast flowing paper sensor. Sensors Actuators B Chem. 418, 136177. 10.1016/j.snb.2024.136177

8. Gabriel, C., Gabriel, S., Corthout, E., 1996. The dielectric properties of biological tissues: I. Literature survey. Phys. Med. Biol. 41, 2231–2249. 10.1088/0031-9155/41/11/001

9. Garcia-Cordero, J.L., Fan, Z.H., 2017. Sessile droplets for chemical and biological assays. Lab Chip 17, 2150–2166. 10.1039/c7lc00366h

10. Giokas, D.L., Tsogas, G.Z., Vlessidis, A.G., 2014. Programming fluid transport in paper-based microfluidic devices using razor-crafted open channels. Anal. Chem. 86, 6202–6207. 10.1021/ac501273v

11. Guo, W., Hansson, J., Van Der Wijngaart, W., 2020. Synthetic Paper Separates Plasma from Whole Blood with Low Protein Loss. Anal. Chem. 92, 6194–6199. 10.1021/acs.analchem.0c01474

12. Hernandez-Perez, R., Fan, Z.H., Garcia-Cordero, J.L., 2016. Evaporation-Driven Bioassays in Suspended Droplets. Anal. Chem. 88, 7312–7317. 10.1021/acs.analchem.6b01657

13. Kar, S., Maiti, T.K., Chakraborty, S., 2015. Capillarity-driven blood plasma separation on paper-based devices. Analyst 140, 6473–6476. 10.1039/c5an00849b

14. Kim, D., Kim, Sejin, Kim, Sanghyo, 2020. An innovative blood plasma separation method for a paper-based analytical device using chitosan functionalization. Analyst 145, 5491–5499. 10.1039/d0an00500b

15. Kontogeorgis, G.M., Kiil, S., 2016. Introduction to Applied Colloid and Surface Chemistry, Introduction to Applied Colloid and Surface Chemistry. 10.1002/9781118881194

16. Lubarda, V.A., Talke, K.A., 2011. Analysis of the equilibrium droplet shape based on an ellipsoidal droplet model. Langmuir 27, 10705–10713. 10.1021/la202077w

17. Modaressi, H., Garnier, G., 2002. Mechanism of wetting and absorption of water droplets on sized paper: Effects of chemical and physical heterogeneity. Langmuir 18, 642–649. 10.1021/la0104931

18. Mukhopadhyay, M., Subramanian, S.G., Durga, K.V., Sarkar, D., DasGupta, S., 2022a. Laser printing based colorimetric paper sensors for glucose and ketone detection: Design, fabrication, and theoretical analysis. Sensors Actuators B Chem. 371, 132599. 10.1016/j.snb.2022.132599

19. Mukhopadhyay, M., Subramanian, S.G., Durga, K.V., Sarkar, D., DasGupta, S., 2022b. Laser printing based colorimetric paper sensors for glucose and ketone detection: Design, fabrication, and theoretical analysis. Sensors Actuators B Chem. 371, 132599. 10.1016/j.snb.2022.132599

20. Nepal, O., Rao, J.P., 2011. Haemolytic effects of hypo-osmotic salt solutions on human erythrocytes. Kathmandu Univ. Med. J. 9, 35–39. 10.3126/kumj.v9i2.6285

21. Nilghaz, A., Shen, W., 2015. Low-cost blood plasma separation method using salt functionalized paper. RSC Adv. 5, 53172–53179. 10.1039/c5ra01468a

22. Park, C., Kim, H.R., Kim, S.K., Jeong, I.K., Pyun, J.C., Park, S., 2019. Three-Dimensional Paper-Based Microfluidic Analytical Devices Integrated with a Plasma Separation Membrane for the Detection of Biomarkers in Whole Blood. ACS Appl. Mater. Interfaces 11, 36428–36434. 10.1021/acsami.9b13644

23. Parsa, M., Harmand, S., Sefiane, K., 2018. Mechanisms of pattern formation from dried sessile drops. Adv. Colloid Interface Sci. 254, 22–47. 10.1016/j.cis.2018.03.007

24. Pitts, K.L., Abu-Mallouh, S., Fenech, M., 2013. Contact angle study of blood dilutions on common microchip materials. J. Mech. Behav. Biomed. Mater. 17, 333–336. 10.1016/j.jmbbm.2012.07.007

25. Pokhrel, P., Jha, S., Giri, B., 2020. Selection of appropriate protein assay method for a paper microfluidics platform. Pract. Lab. Med. 21, e00166. 10.1016/j.plabm.2020.e00166

26. Songjaroen, T., Dungchai, W., Chailapakul, O., Henry, C.S., Laiwattanapaisal, W., 2012. Blood separation on microfluidic paper-based analytical devices. Lab Chip 12, 3392–3398. 10.1039/c2lc21299d

27. Toyoda, T., Isobe, K., Tsujino, T., Koyata, Y., Ohyagi, F., Watanabe, T., Nakamura, M., Kitamura, Y., Okudera, H., Nakata, K., Kawase, T., 2018. Direct activation of platelets by addition of CaCl2 leads coagulation of platelet-rich plasma. Int. J. Implant Dent. 4. 10.1186/s40729-018-0134-6

28. Trantum, J.R., Baglia, M.L., Eagleton, Z.E., Mernaugh, R.L., Haselton, F.R., 2014. Biosensor design based on Marangoni flow in an evaporating drop. Lab Chip 14, 315–324. 10.1039/c3lc50991e

29. Yang, M., Chen, D., Hu, J., Zheng, X., Lin, Z.J., Zhu, H., 2022. The application of coffee-ring effect in analytical chemistry. TrAC - Trends Anal. Chem. 157, 116752. 10.1016/j.trac.2022.116752

30. Yang, X., Forouzan, O., Brown, T.P., Shevkoplyas, S.S., 2012. Integrated separation of blood plasma from whole blood for microfluidic paper-based analytical devices. Lab Chip 12, 274–280. 10.1039/c1lc20803a

