## Supplemental File for "A Paper-Droplet Hybrid System for Efficient Blood Plasma Separation in Multiplex Diagnostics"

**S1**. *Substate preparation for Droplet Residing:*

The preparation of a hydrophobic substate is essential for residing and maintaining stability of the droplet during blood plasma separation. Here, a hydrophobic paper substate is prepared using simplified Laser Printing technique for low-cost and easy implementation in point-of-care application. First, grade-1 filter paper is adhered to a regular A4 size printing paper using glue (Fevistick non-toxic adhesive), applied only at the edges of the filter paper for easy removal after printing. The filter paper attached to the A4 sheet is then inserted into an HP Laser Pro 400 series printer and printed using regular printing process. A predesigned black region, prepared using Inkscape (version 1.3.1) was printed on the attached filter paper. Once the filter paper is printed black, it is removed from the host A4 sheet. The filter paper is then heated for 1 hour at 180°C using a hotplate (Tarson Hot top Digital MC-02) allowing the ink to percolate into the paper matrix and form a hydrophobic barrier (Mukhopadhyay et al., 2022). The schematic of the entire substrate preparation process is shown in Fig. S1

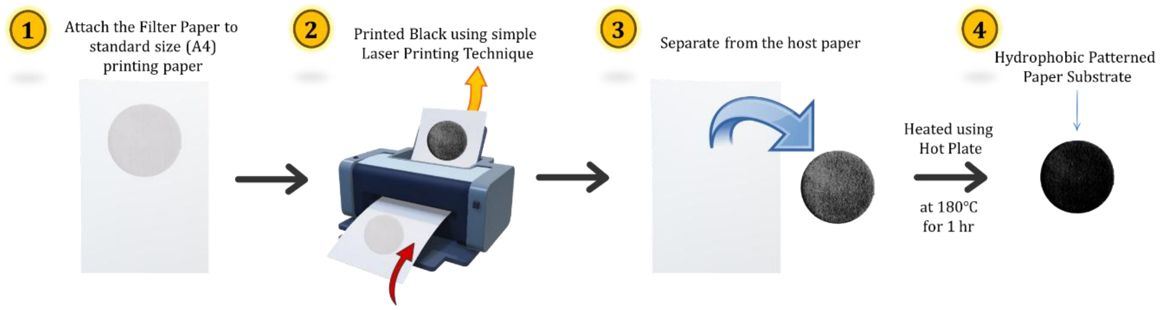

**Fig. S1** Schematic of the process for making hydrophobic patterning on porous paper matrix for stable residing of blood and salt solution droplet mixture.

**S2**. *Dimension Optimization of Paper Channels:*

A paper strip functioning as paper microfluidic platform (μPAD) was designed using Inkscape (version 1.3.1). The designs were directly printed on grade-1 filter paper following steps 1, 2 and 3 shown in Fig. S1 and sheared into individual paper devices manually. The dimensions of the device and each zone (separation/test zone, buffer zone and detection zone) are optimized to maximize separation bandwidth of plasma from whole blood, where separation bandwidth $(\Delta l_{sep.}) = Max.$ plasma traverse length $(l_{plasma}) - Max.$ RBC traverse length $(l_{RBC})$; and subsequent multiplexing of plasma biomarkers. The length traversed by red blood cells (RBCs) and plasma, as well as the difference between them $(\Delta l_{sep.})$, are tabulated in Table S1.

**Table S1:** Length traversed by red blood cells (RBCs), plasma and separation bandwidth for varied salt dilution.

| $\left(\frac{V_{blood\ sample}}{V_{salt\ solution}}\right)$ | **Channel Width** | | | | | | | | |
|---|---|---|---|---|---|---|---|---|---|
| | **4 mm** | | | **8 mm** | | | **10 mm** | | |
| | RBC (cm) | Plasma (cm) | Bandwidth (cm) | RBC (cm) | Plasma (cm) | Bandwidth (cm) | RBC (cm) | Plasma (cm) | Bandwidth (cm) |
| 10 μL (Blood) + 0 μL ($CaCl_2$) | 0.4 $\pm$ 0.05 | NA | NA | 0.36 $\pm$ 0.05 | NA | NA | 0.46 $\pm$ 0.11 | NA | NA |
| 9 μL (Blood) + 1 μL ($CaCl_2$) | 0.76 $\pm$ 0.15 | 0.9 $\pm$ 0.1 | 0.13 $\pm$ 0.05 | 0.73 $\pm$ 0.05 | 0.90 $\pm$ 0.10 | 0.16 $\pm$ 0.05 | 0.30 $\pm$ 0.00 | 0.60 $\pm$ 0.00 | 0.30 $\pm$ 0.00 |
| 8 μL (Blood) + 2 μL ($CaCl_2$) | 0.5 $\pm$ 0.1 | 1.16 $\pm$ 0.05 | 0.66 $\pm$ 0.15 | 0.53 $\pm$ 0.11 | 1.16 $\pm$ 0.05 | 0.63 $\pm$ 0.15 | 0.36 $\pm$ 0.05 | 0.73 $\pm$ 0.23 | 0.36 $\pm$ 0.20 |
| 7 μL (Blood) + 3 μL ($CaCl_2$) | 0.5 $\pm$ 0.05 | 1.23 $\pm$ 0.05 | 0.73 $\pm$ 0.05 | 0.46 $\pm$ 0.05 | 1.26 $\pm$ 0.05 | 0.80 $\pm$ 0.10 | 0.43 $\pm$ 0.15 | 0.86 $\pm$ 0.30 | 0.43 $\pm$ 0.2 |
| 6 μL (Blood) + 4 μL ($CaCl_2$) | 0.46 $\pm$ 0.05 | 1.33 $\pm$ 0.05 | 0.86 $\pm$ 0.05 | 0.56 $\pm$ 0.05 | 1.36 $\pm$ 0.05 | 0.80 $\pm$ 0.10 | 0.23 $\pm$ 0.05 | 0.7 $\pm$ 0.00 | 0.46 $\pm$ 0.05 |
| 5 μL (Blood) + 5 μL ($CaCl_2$) | 0.5 $\pm$ 0.05 | 1.4 $\pm$ 0.00 | 0.9 $\pm$ 0.00 | 0.53 $\pm$ 0.05 | 1.36 $\pm$ 0.05 | 0.83 $\pm$ 0.05 | 0.36 $\pm$ 0.05 | 0.93 $\pm$ 0.11 | 0.56 $\pm$ 0.15 |

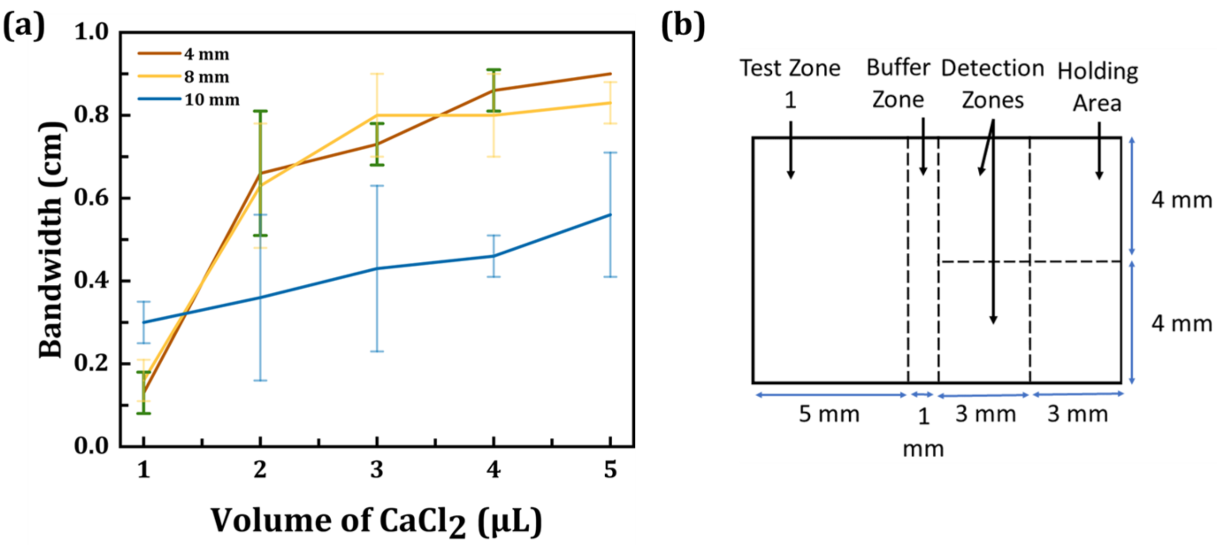

**Fig. S2** **(a)** Plot of variation in separation bandwidth (cm) of plasma with increase in volume of $CaCl_2$ salt solution (μL) **(b)** Optimised channel dimension for maximum bandwidth of plasma separation from whole blood and subsequent multiplexing of plasma biomarkers.

With increase in channel width, the separation bandwidth decreases. For 8 mm channel width however, the flow was comparable to 4 mm channel and therefore was considered for further experimentation for easy handling without compromising separation. All the wicking experiments were performed for different volume of salt solution. Although with increase in volume of salt solution, the separation bandwidth increases, it simultaneously dilutes the blood sample for subsequent analysis. This can compromise with the accuracy and increase the limit-of-detection (LOD) which is not desirable. Also, with about 1: 1 dilution, the separation bandwidth starts to saturate as shown in Fig. S2(a). Therefore, the dilution ratio is restricted to 1:1 blood sample to salt solution. The final device design with all the regions marked and dimension specified is shown in Fig. S2(b).

#### **S3**. *Blood Plasma Separation preliminary testing:*

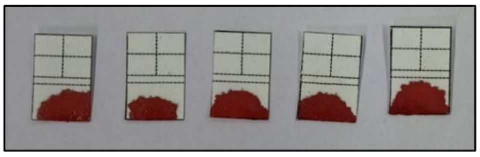

**Fig. S3** Control Blood plasma separation testing without any salt solution, using only whole blood. Only large blood patches were observed as such and no separation was recorded without salts.

**Table S2:** Blood plasma separation performed by storing 5 healthy blood samples* for 5 days, each repeated 5 times per day (5×5×5=125 tests). Two different samples with hematocrit, Hct = 34.1% and 45.8% are shown here, with consistent separation.

| **Day No.** | **Hct = 34.1 %** | **Hct = 45.8 %** |
|---|---|---|
| **1.** | 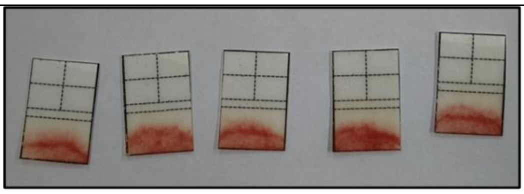 | 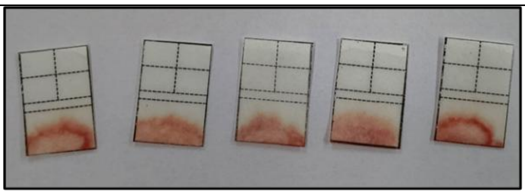 |
| **2.** | 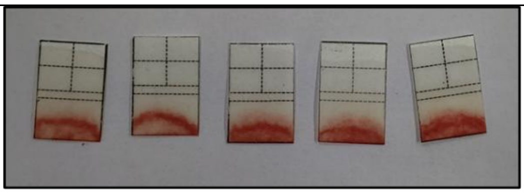 | 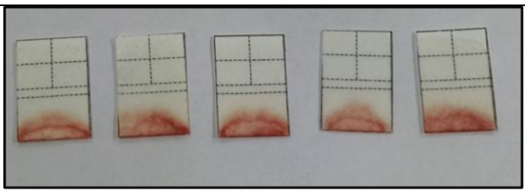 |

| | | |
|---|---|---|
| **3.** | 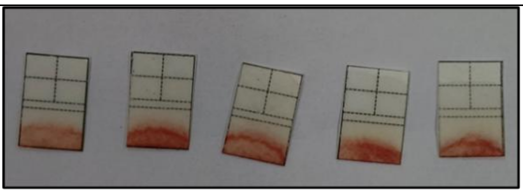 | 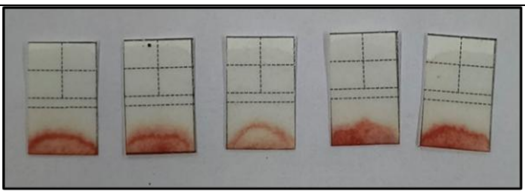 |
| **4.** | 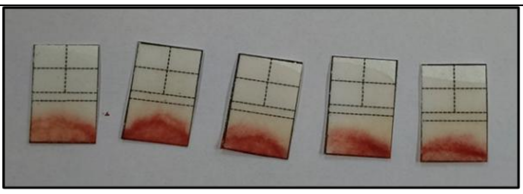 | 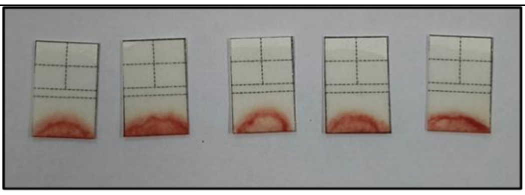 |
| **5.** | 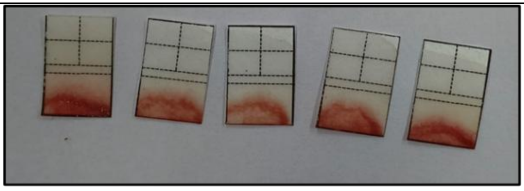 | 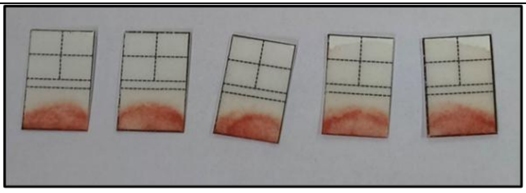 |

*Blood is stored in the refrigerator with a temperature between $2 - 6\,^{o}C$ to prevent hemolysis, growth of bacterial contamination. Before use every day, the blood samples were taken out on an average 30 min before analysis to allow the blood to reach normal temperature.

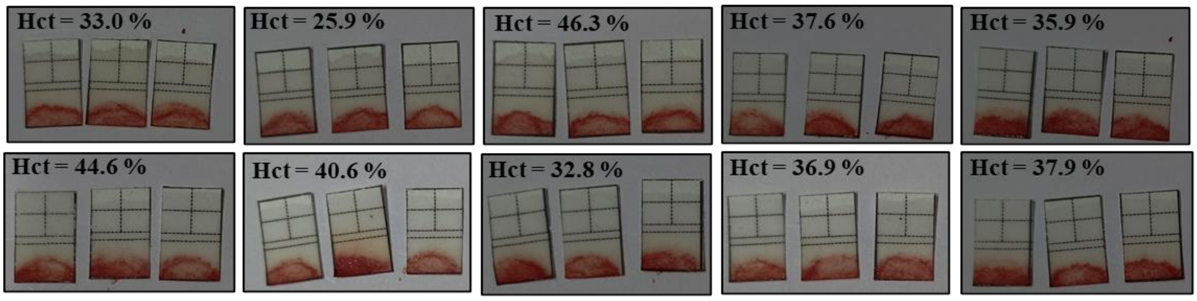

**Fig. S4** Blood plasma separation performed using 35 healthy blood samples (35×3=105 tests) with a wide hematocrit spectrum. 10 samples are shown here.

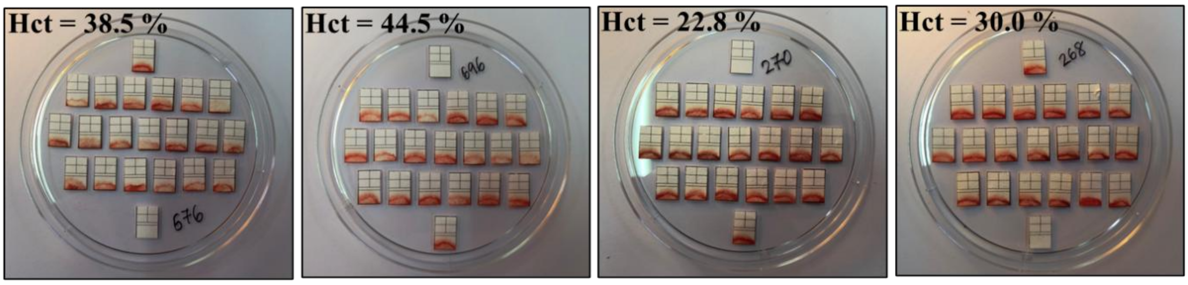

**Fig. S5** Reproducibility test performed by repeating separation with 10 samples, each for 20 times (10×20=200 tests). Four samples are shown here.

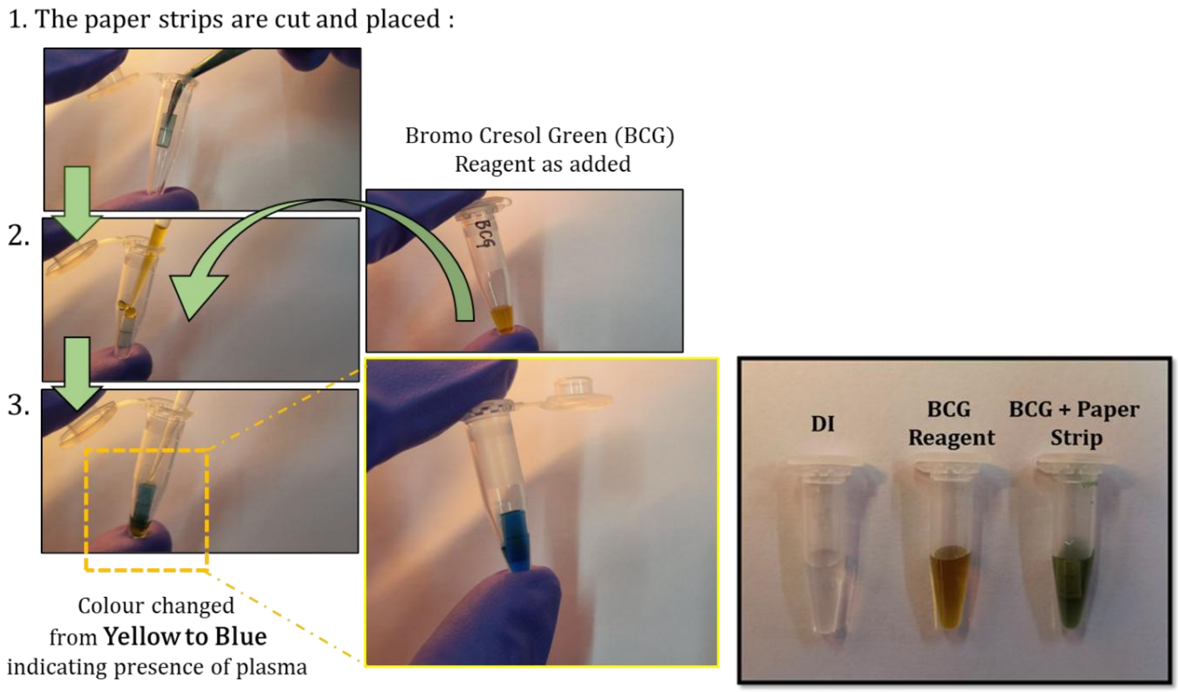

**Fig. S6**** Preliminary testing of separated plasma is performed using Bromocresol Green (BCG) solution. In three 0.5 mL standard centrifuge tubes, 200 μL of DI water is taken in one and BCG solution in taken in rest two. The detection zone was cut from the paper strip and dipped in one of the BCG solutions. The centrifuge tube was kept in a water bath and sonicated for 15 min. The change in colour from yellow to blue indicates the presence of plasma in the detection zones.

**Here, we have used Labogen's Bromocresol Green indicator solution (0.04%), pH: 3.6-5.4, Yellow to Blue which is standard reagent for labroatory albumin detection.

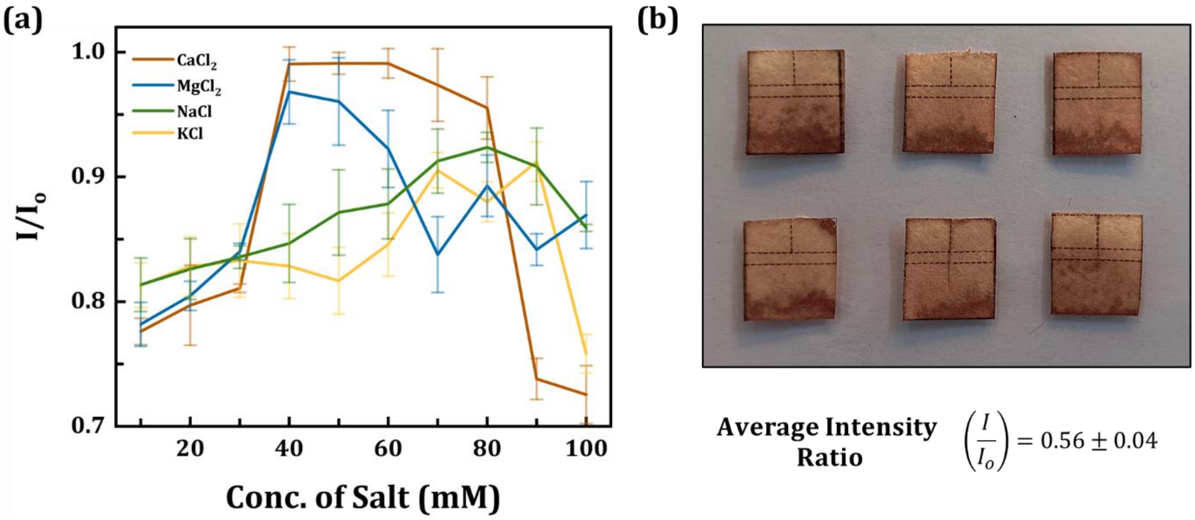

**Fig. S7** **(a)** Intensity ratios $\left(\frac{I}{I_o}\right)$ of the detection zones for varied concentration of 4 different salts: The images were captured using Optical Microscope (Leica DM6000M) with 5× objective lens and analysed using ImageJ (mean $\pm$ SD, $n = 5$) **(b)** Actual device of completely hemolysed blood obtained by mixing DI & used for blood plasma separation. Average intensity ratio obtained from the 12 (6 devices × 2 detection zones) detection zone is used as the control ($IR_{control}$ ) to evaluate quality of plasma obtained ($\eta$)

### **S4**. *Effects of Salts on Blood Plasma Separation:*

**Table S3:** Calculated values of plasma yield quality ($\eta$) using eq. (1) at the detection zone of the paper channel

| Conc. (mM) | 10 | 20 | 30 | 40 | 50 | 60 | 70 | 80 | 90 | 100 |
|---|---|---|---|---|---|---|---|---|---|---|
| $\boldsymbol{\eta}$ | 0.479 $\pm$ 0.06 | 0.528 $\pm$ 0.10 | 0.560 $\pm$ 0.03 | 0.977 $\pm$ 0.03 | 0.979 $\pm$ 0.02 | 0.978 $\pm$ 0.07 | 0.938 $\pm$ 0.07 | 0.895 $\pm$ 0.06 | 0.391 $\pm$ 0.08 | 0.361 $\pm$ 0.09 |

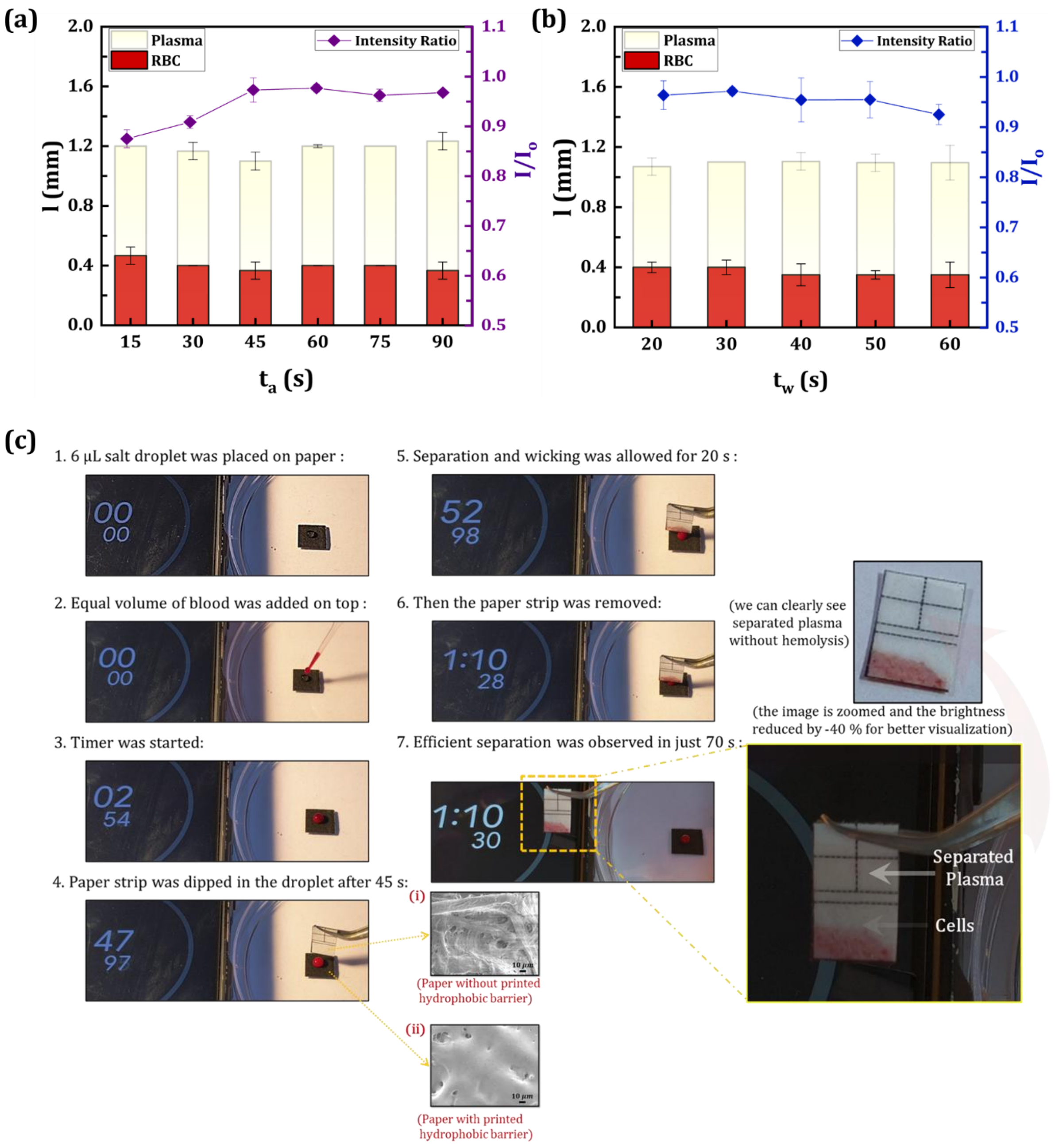

**Fig. S8** **(a)** & **(b)** Distance, $l$(mm) traversed by RBCs & separated plasma; and separation extent at detection zone for varied cell aggregation time ($t_a$) and paper strip wicking time ($t_w$); (mean $\pm$ SD, $n = 5$). **(c)** Steps in plasma separation using blood drop, salt solution (40mM $CaCl_2$) and paper. Time was recorded in smartphone (~70 s) and image of final device with separated plasma is magnified and highlighted separately.

### S5. *Separation and Detection Zone Optimization:*

Here, for separation zone optimization we first observed what is the minimum length the cells are traversing through the paper strip. For that we considered randomly 10 devices from Table S2 with observable minimum cell transport in paper and obtained the fractional area coverage using self-written MATLAB code. The area covered by the cell w.r.t the entire strip is given in Table S4. Since, the cells cover an observable minimum area of around ≈ 27 % fraction, we set the separation region around twice this area ≈ 50 % i.e. 6mm length. We observed that in almost all the devices the cells could not traverse any further that 50 % of the area, as shown in Fig S2(b), and therefore was considered for final design.

**Table S4:** The fractional area coverage of the cells obtained for 10 devices from Table S2. The fractional area coverage is defined as area covered by cells w.r.t entire paper strip.

| | | | | | |
|---|---|---|---|---|---|
| Device Image | 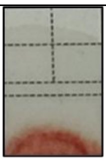 | 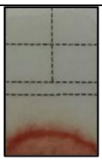 | 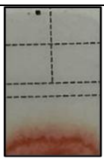 | 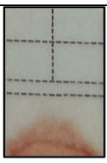 | 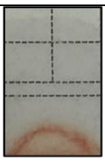 |
| Fractional Area Coverage | 25.89 % | 36.24 % | 32.12 % | 22.24 % | 22.47 % |
| Device Image | 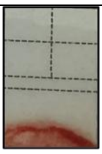 | 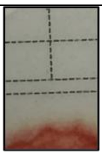 | 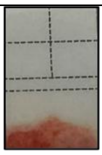 | 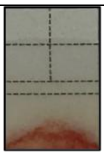 | 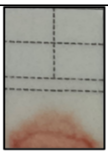 |
| Fractional Area Coverage | 27.11 % | 29.82 % | 28.93 % | 24.73 % | 23.57 % |
| **Total Average Minimum Area Coverage ≈ 27 ± 5 %** | | | | | |

**S6.** *Multi-analyte detection following BPS:*

The glucose and albumin assay are performed using previously established glucose and albumin assay method in paper analytical device (Kim et al., 2020), (Nilghaz and Shen, 2015), (Yang et al., 2012), (Songjaroen et al., 2012), (Ardakani and Hemmateenejad, 2023), (Pokhrel et al., 2020). The detection zones were stained with 2 $\mu$L of reagent each. All the reagents used however are optimised according to the current assay to obtain low limit-of-detection (LOD) simultaneously incorporating analyte range of physiological importance. The science behind each assay and reagents used are presented below:

*S5.1. Glucose Assay:*

We used an enzymatic approach for glucose detection, as it offers more specificity compared to other approaches, because of the reduced presence of interfering substances and more stable reagents. Furthermore, this method is advantageous owing to its simplicity, since it can be accomplished in a single step. We have used the Glucose Oxidase-Peroxidase (GODPOD) technique. The methodology and process for the glucose test have been well established in prior research. Dextrose was used as a reference standard for calibration investigations. At room temperature, the enzymatic interaction between glucose and glucose oxidase (GOx) in the presence of oxygen results in the production of glucono-delta-lactone and hydrogen peroxide. In the presence of Horseradish peroxidase (HRP), the hydrogen peroxide by-product combines with potassium iodide, causing the oxidation of iodide to iodine and producing a characteristic golden yellow colouring. The intensity of this developed colour will be directly proportional the concentration of glucose in the plasma.

$$Glucose + O_2 \xrightarrow{GO_X} Glucono - delta - lactone + H_2O_2$$

$$H_2O_2 + 2I^- \xrightarrow{HRP} H_2O + I_2$$

*Reagent:*

The glucose reagent comprises two solutions: (1) 50 mM KI and 0.3 M Trehalose in 10× Phosphate Buffer at pH 7.4, mixed with Triton X-100 at a ratio of 10:1 (2) 120 units/mL HRP and 120 units/mL GOx mixed in 1:1 ratio. The reagent consists of an equal proportion of both reagents, resulting in the formation of the glucose detection reagent. 2 µL of the glucose detection reagent was spotted to dedicated section of the detection zone and allowed to dry. Dextrose concentrations ranging from 30 to 150 mg/dL were prepared and stored in a vial for calibration studies from 200 mg/dL stock solution.

*S5.2. Albumin Assay:*

We used Bromocresol Green method for albumin detection in plasma. For baseline calibration studies, Human Serum Albumin (HSA) is considered a gold standard for protein determination. In this method the specific binding of bromocresol green (BCG), an anionic dye, and the protein at acid pH produce a colour change of the indicator from yellow –green to green –blue. The intensity of the colour formed is proportional to the concentration of albumin in the sample.

$$BCG + Albumin \xrightarrow{pH\ 4.3} BCG - Albumin\ Complex$$

*Reagent:*

The albumin reagent consists of an equal proportion of 1.7 mg/mL of BCG mixed with 30 wt% PEG-10000, spotted on the detection zone of the paper strip. HSA concentrations ranging from 0 to 10 g/dL were prepared and stored in a vial for calibration studies from 30 mg/mL stock solution. The reagent reacts with BCG to yield Albumin-BCG complex (Teal coloration).

**S7.** *Statistics:*

- Statistics for the Linear Regression Model:

Since the slopes are numerically very small; 0.0012 for glucose and 0.0177 for albumin, we performed the test of significance for both the regression studies. This provides an objective way to evaluate whether the small slope is meaningful in the context of the data and model. There are two tests of significance for regression analysis i.e. ANOVA and t-statistics. For a small slope, ANOVA helps in determining if the regression model significantly improves the prediction compared to no model. Similarly, the t-statistics can tell whether the smaller slope is likely real or just due to random variation in the data.

For a regression equation of: $y = mx + c$

1. **ANOVA:** (Test of significance for the entire regression)

$Null\ Hypothesis$ $H_0 : m = 0$ ($x$ no regression relationship exists)
$Alternate\ Hypothesis$ $H_1 : m \neq 0$ (regression relationship exists)

2. **t-statistics:** (Test of significance for the slope)

$Null\ Hypothesis$ $H_0 : m = 0$ ($x$ has no influence on $y$)

$Alternate\ Hypothesis\ \ H_1 : m \neq 0$

**(a)**

| ANOVA | | | | | | | | |
|---|---|---|---|---|---|---|---|---|
| | *df* | *SS* | *MS* | *F* | *Significance F* | | | |
| Regression | 1 | 0.010531984 | 0.010531984 | 150.6506992 | 1.78174E-05 | | | |
| Residual | 6 | 0.00041946 | 6.991E-05 | | | | | |
| Total | 7 | 0.010951444 | | | | | | |
| | *Coefficients* | *Standard Error* | *t Stat* | *P-value* | *Lower 95%* | *Upper 95%* | *Lower 95.0%* | *Upper 95.0%* |
| Intercept | 0.98053973 | 0.008487452 | 115.5281561 | 2.83573E-11 | 0.959771682 | 1.001307777 | 0.959771682 | 1.001307777 |
| X Variable 1 | -0.105569677 | 0.008601092 | -12.27398465 | 1.78174E-05 | -0.126615792 | -0.084523562 | -0.126615792 | -0.084523562 |

**(b)**

| ANOVA | | | | | | | | |
|---|---|---|---|---|---|---|---|---|
| | *df* | *SS* | *MS* | *F* | *Significance F* | | | |
| Regression | 1 | 0.016237952 | 0.016237952 | 356.9873039 | 2.89233E-07 | | | |
| Residual | 7 | 0.000318403 | 4.54861E-05 | | | | | |
| Total | 8 | 0.016556355 | | | | | | |
| | *Coefficients* | *Standard Error* | *t Stat* | *P-value* | *Lower 95%* | *Upper 95%* | *Lower 95.0%* | *Upper 95.0%* |
| Intercept | 0.506782039 | 0.004899647 | 103.4323559 | 2.08112E-12 | 0.495196215 | 0.518367863 | 0.495196215 | 0.518367863 |
| X Variable 1 | 0.001645091 | 8.7069E-05 | 18.89410765 | 2.89233E-07 | 0.001439206 | 0.001850977 | 0.001439206 | 0.001850977 |

**Fig. S9** **(a)** & **(b)** ANOVA and t-statistics data for glucose and albumin calibration linear regression model for the major concentrations.

- Student T-test for 25 random concentrations:

This test is used to check if there is any difference exist between two independent sample set. It is generally used when the sample size is $\leq 30$.

$Null\ Hypothesis$ $H_0$ : There is no difference between both the data set

$Alternate\ Hypothesis$ $H_1$ : There exist a difference between both the data set

**(a)**

| t-Test: Two-Sample Assuming Unequal Variances | | |
|---|---|---|
| | *Variable 1* | *Variable 2* |
| Mean | 0.886002815 | 0.8874056 |
| Variance | 0.001083373 | 0.001105158 |
| Observations | 25 | 25 |
| Hypothesized Mean Difference | 0 | |
| df | 48 | |
| t Stat | -0.149928654 | |
| P(T<=t) one-tail | 0.440724566 | |
| t Critical one-tail | 1.677224196 | |
| P(T<=t) two-tail | 0.881449133 | |
| t Critical two-tail | 2.010634758 | |

**(b)**

| t-Test: Two-Sample Assuming Unequal Variances | | |
|---|---|---|
| | *Variable 1* | *Variable 2* |
| Mean | 0.630200616 | 0.6354912 |
| Variance | 0.000814531 | 0.001015007 |
| Observations | 25 | 25 |
| Hypothesized Mean Difference | 0 | |
| df | 47 | |
| t Stat | -0.618447534 | |
| P(T<=t) one-tail | 0.269633077 | |
| t Critical one-tail | 1.677926722 | |
| P(T<=t) two-tail | 0.539266154 | |
| t Critical two-tail | 2.011740514 | |

**Fig. S10** **(a)** & **(b)** Student T-test data for 25 random data points for glucose in the 40-130 mg/dL and for albumin in the 1-9 g/dL linearly fit calibration range.

### **S8.** *Healthy Blood Specifications:*

Individuals, with $50 \pm 20$ kg of body-weight, $73 \pm 4$ beats/min of resting heart rate, $120.5 \pm 7$ mmHg of systolic and $84 \pm 3$ mm Hg of diastolic blood pressure including non-smokers and devoid of any disease and disorder are included in the analysis.

**Table S5:** The specific blood parameters and their ranges obtained from Automatic Haematology Analyser (Sysmex, KX-21), for the healthy individuals, are listed below.

| **SR. NO** | **Parameter** | **Range** | **Unit** |
|---|---|---|---|
| 1 | WBC | 3.5-9.5 | $10^3$/μL |
| 2 | Neu% | 40-75 | % |
| 3 | Lym% | 20-50 | % |
| 4 | Mon% | 3-10 | % |
| 5 | Eos% | 0.4-8 | % |
| 6 | Bas% | 0-1 | % |
| 7 | Neu# | 1.8-6.3 | $10^3$/μL |
| 8 | Lym# | 1.1-3.2 | $10^3$/μL |
| 9 | Mon# | 0.1-0.6 | $10^3$/μL |
| 10 | Eos# | 0.02-0.52 | $10^3$/μL |
| 11 | Bas# | 0-0.06 | $10^3$/μL |
| 12 | ALY# | 0-0.2 | $10^3$/μL |
| 13 | ALY% | 0-2 | % |
| 14 | LIC# | 0-0.02 | $10^3$/μL |
| 15 | LIC% | 0-0.1 | % |
| 16 | NRBC | 0.00 | $10^3$/μL |

| **SR. NO** | **Parameter** | **Range** | **Unit** |
|---|---|---|---|
| 17 | NRBC% | 0.00 | % |
| 18 | RBC | 3.8-5.8 | $10^6$/μL |
| 19 | HGB | 11-17 | g/dL |
| 20 | HCT | 35-50 | % |
| 21 | MCV | 82-100 | fL |
| 22 | MCH | 27-34 | Pg |
| 23 | MCHC | 31-35 | g/dL |
| 24 | RDW-CV | 11-16 | % |
| 25 | RDW-SD | 35-56 | fL |
| 26 | PLT | 125-350 | $10^3$/μL |
| 27 | MPV | 6.5-12 | fL |
| 28 | PDW-SD | 9-17 | fL |
| 29 | PDW-CV | 10-17.9 | % |
| 30 | PCT | 0.1-0.2 | % |
| 31 | P-LCR | 11-45 | % |
| 32 | P-LCC | 30-90 | $10^3$/μL |

### S9. *Experimentation with Real Blood Samples:*

**Table S6:** Preliminary testing of blood plasma separation and simultaneously detecting glucose and albumin levels in 5 healthy blood samples. Each sample was performed in triplicate and validated with the 95% prediction band.

| | | | | | | |
|---|---|---|---|---|---|---|
| **Sample 1** | **Hematocrit (%)** | **Glucose (mg/dL)** | **IR (1)** | **IR (2)** | **IR (3)** | **% RSD** |
| | 44.6 | 100 | 0.914974 | 0.931785 | 0.965648 | 2.7 |
| | **Total Protein (g/dL)** | **Albumin (g/dL)** | **IR (1)** | **IR (2)** | **IR (3)** | **% RSD** |
| | 7.2 | 4.2 | 0.660155 | 0.664097 | 0.668913 | 0.6 |
| **Sample 2** | **Hematocrit (%)** | **Glucose (mg/dL)** | **IR (1)** | **IR (2)** | **IR (3)** | **% RSD** |
| | 39 | 108 | 0.856465 | 0.85772 | 0.882595 | 1.7 |
| | **Total Protein (g/dL)** | **Albumin (g/dL)** | **IR (1)** | **IR (2)** | **IR (3)** | **% RSD** |
| | 7 | 4.3 | 0.651849 | 0.664041 | 0.609826 | 4.4 |
| **Sample 3** | **Hematocrit (%)** | **Glucose (mg/dL)** | **IR (1)** | **IR (2)** | **IR (3)** | **% RSD** |
| | 34.6 | 101 | 0.866356 | 0.853037 | 0.862581 | 0.8 |
| | **Total Protein (g/dL)** | **Albumin (g/dL)** | **IR (1)** | **IR (2)** | **IR (3)** | **% RSD** |
| | 6.8 | 4.1 | 0.572795 | 0.578832 | 0.577025 | 0.5 |
| **Sample 4** | **Hematocrit (%)** | **Glucose (mg/dL)** | **IR (1)** | **IR (2)** | **IR (3)** | **% RSD** |
| | 37.9 | 96 | 0.915176 | 0.936726 | 0.913936 | 1.4 |
| | **Total Protein (g/dL)** | **Albumin (g/dL)** | **IR (1)** | **IR (2)** | **IR (3)** | **% RSD** |
| | 6 | 3.8 | 0.627163 | 0.604229 | 0.605821 | 2 |
| **Sample 5** | **Hematocrit (%)** | **Glucose (mg/dL)** | **IR (1)** | **IR (2)** | **IR (3)** | **% RSD** |
| | 38 | 89 | 0.860652 | 0.85614 | 0.856097 | 0.3 |
| | **Total Protein (g/dL)** | **Albumin (g/dL)** | **IR (1)** | **IR (2)** | **IR (3)** | **% RSD** |
| | 7 | 3.6 | 0.602354 | 0.646265 | 0.667384 | 5.1 |

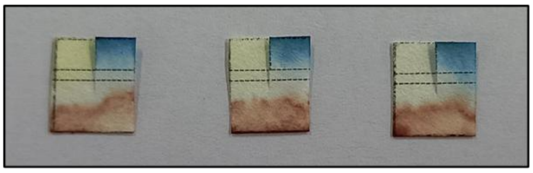

**Fig. S11** Highlighted images of the paper device in Table S6 above performed in triplicate

**Table S7:** Testing of whole blood for blood plasma separation and simultaneously detecting glucose and albumin levels in 30 healthy, 30 diluted and 30 spiked blood samples. All the samples and corresponding data are obtained from B. C. Roy technology Hospital, IIT Kharagpur. The hematocrit (Hct) data was obtained from Automatic Haematology Analyser (Sysmex, KX-21) and the glucose (Glu.), albumin (Alb.) and total protein (TP) data was obtained from Automated Biochemistry Analyzer (DiaSYS 400)

| SR. No. | Hct (%) | TP (g/dL) | | Unmodified Whole Blood Samples | | | Diluted Whole Blood Samples * | | | Spiked Whole Blood Samples ** | | |
|---|---|---|---|---|---|---|---|---|---|---|---|---|
| | | | | Sample No. | Glu. (mg/dL) | Alb. (g/dL) | Sample No. | Glu. (mg/dL) | Alb. (g/dL) | Sample No. | Glu. (mg/dL) | Alb. (g/dL) |
| 1 | 41.1 | 7.9 | | **1** | 107 | 4.9 | **31** | 70 | 3.2 | **61** | 70 | 7.3 |
| 2 | 36.9 | 6.5 | | **2** | 92 | 4 | **32** | 62 | 2.7 | **62** | 62 | 6.5 |
| 3 | 34.1 | 5.9 | | **3** | 113 | 3.5 | **33** | 78 | 2.4 | **63** | 78 | 6.0 |
| 4 | 34.8 | 7.7 | | **4** | 140 | 4.4 | **34** | 97 | 3.0 | **64** | 97 | 6.7 |
| 5 | 44 | 7.7 | | **5** | 82 | 4.5 | **35** | 52 | 2.8 | **65** | 52 | 7.1 |
| 6 | 44.5 | 7.2 | | **6** | 143 | 4.7 | **36** | 91 | 3.0 | **66** | 91 | 7.3 |
| 7 | 45.8 | 7 | | **7** | 125 | 4.6 | **37** | 78 | 2.9 | **67** | 78 | 7.3 |
| 8 | 30.7 | 6.4 | | **8** | 136 | 4 | **38** | 78 | 2.8 | **68** | 96 | 6.3 |
| 9 | 37.6 | 6.6 | | **9** | 159 | 4.1 | **39** | 108 | 2.7 | **69** | 108 | 6.6 |
| 10 | 37.6 | 6.7 | | **10** | 178 | 4.1 | **40** | 120 | 2.7 | **70** | 120 | 6.6 |
| 11 | 38 | 7 | | **11** | 89 | 3.6 | **41** | 60 | 3.1 | **71** | 60 | 6.9 |
| 12 | 40.6 | 6.9 | | **12** | 98 | 4.6 | **42** | 65 | 3.0 | **72** | 65 | 7.0 |
| 13 | 33.6 | 6 | | **13** | 91 | 4 | **43** | 63 | 2.7 | **73** | 63 | 6.4 |
| 14 | 38.8 | 5.9 | | **14** | 118 | 3.6 | **44** | 79 | 2.4 | **74** | 79 | 6.3 |
| 15 | 39.4 | 6.6 | | **15** | 238 | 4.1 | **45** | 159 | 2.7 | **75** | 159 | 6.7 |
| 16 | 46.3 | 6.9 | | **16** | 95 | 4.5 | **46** | 59 | 2.8 | **76** | 59 | 7.2 |

| | | | | | | | | | | | | |
|---|---|---|---|---|---|---|---|---|---|---|---|---|
| 17 | 44.6 | 7.2 | | **17** | 100 | 4.2 | **47** | 63 | 2.6 | **77** | 63 | 7.0 |
| 18 | 37.9 | 6 | | **18** | 96 | 3.8 | **48** | 65 | 2.5 | **78** | 65 | 6.4 |
| 19 | 38 | 7.6 | | **19** | 346 | 4.3 | **49** | 234 | 2.9 | **79** | 234 | 6.3 |
| 20 | 39.9 | 7.2 | | **20** | 137 | 4.2 | **50** | 91 | 2.8 | **80** | 91 | 6.7 |
| 21 | 32.8 | 7 | | **21** | 108 | 4.2 | **51** | 75 | 2.9 | **81** | 75 | 6.5 |
| 22 | 34.6 | 6.8 | | **22** | 101 | 4.1 | **52** | 70 | 2.8 | **82** | 70 | 6.5 |
| 23 | 31.2 | 6.9 | | **23** | 85 | 4.2 | **53** | 60 | 2.9 | **83** | 60 | 6.4 |
| 24 | 25.9 | 5.8 | | **24** | 101 | 3.2 | **54** | 73 | 2.3 | **84** | 73 | 5.5 |
| 25 | 34.6 | 6.1 | | **25** | 98 | 4.1 | **55** | 68 | 2.8 | **85** | 68 | 6.5 |
| 26 | 35.5 | 6.4 | | **26** | 87 | 4.1 | **56** | 60 | 2.8 | **86** | 60 | 6.5 |
| 27 | 38.6 | 6.1 | | **27** | 291 | 4.4 | **57** | 196 | 2.9 | **87** | 196 | 6.8 |
| 28 | 35.9 | 6.7 | | **28** | 86 | 4 | **58** | 59 | 2.7 | **88** | 59 | 6.4 |
| 29 | 39 | 7 | | **29** | 108 | 4.3 | **59** | 72 | 2.8 | **89** | 72 | 6.8 |
| 30 | 33 | 6.6 | | **30** | 114 | 3.9 | **60** | 79 | 2.7 | **90** | 79 | 6.3 |

* & **, both the diluted and spiked whole blood samples are prepared from the respective unmodified whole blood sample present in the same row. Modification was done by first separating 100 μL of blood sample and centrifuged at 3000 rpm for 10 min. For dilution, 20 μL of supernatant plasma was replaced with 10 × PBS. For spike, 20 μL of supernatant plasma was replaced with 1.2 g/dL HSA solution in 10 × PBS. The samples are mixed and used.

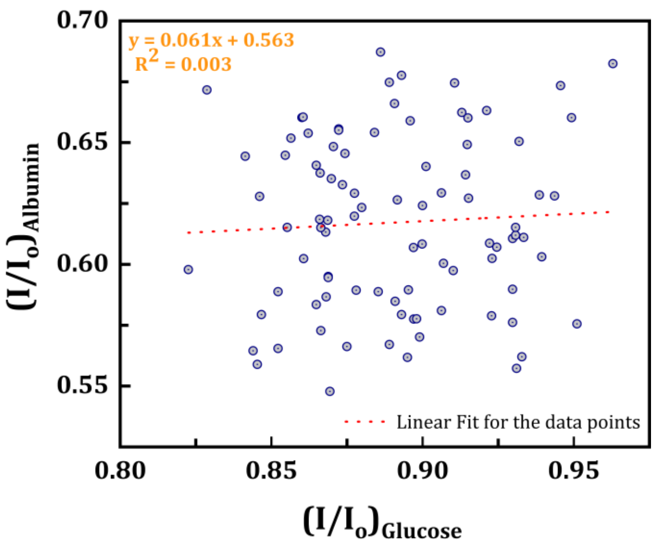

**Fig. S12** Scatter point plot of intensity ratio of albumin, $\left(\frac{I}{I_o}\right)_{Albumin}$ vs. intensity ratio of glucose, $\left(\frac{I}{I_o}\right)_{Glucose}$ of each test, to study the cross-dependence of colour developed between glucose and albumin assay in a single strip. The distribution appears random and no specific dependence could be identified.

### S10. *MATLAB GPS Application:*

**Table S8:** Comparison of Intensity Ratio values obtained from MATLAB GPS Application and that obtained from ImageJ for 5 samples tabulated in Table S7 and highlighted in green

| SR. No. | **IR obtained from MATLAB GPS App.** (for Glucose) | **IR obtained from ImageJ** (for Glucose) | **IR obtained from MATLAB GPS App.** (for Albumin) | **IR obtained from ImageJ** (for Albumin) |
|---|---|---|---|---|
| 1 | 0.864 | 0.866 | 0.583 | 0.580 |
| 2 | 0.906 | 0.902 | 0.581 | 0.580 |
| 3 | 0.922 | 0.921 | 0.608 | 0.609 |
| 4 | 0.856 | 0.856 | 0.651 | 0.654 |
| 5 | 0.864 | 0.863 | 0.640 | 0.637 |

The developed MATLAB GPS Application aim to streamline the quantitative estimation of test results and minimize error. Since the application relies on image processing, using quality images is essential. The application calculates the intensity ratio of two images i.e. the image of glucose and protein region with that of a blank image for reference, so all images must be captured under consistent lighting and camera settings. The image that are used in Supplementary Video S2 were captured using Moto Edge 40 Neo smartphone camera in manual mode with consistent setting. The used images are attached below. The quality and consistency of the obtained image is very much crucial for accurate results, as any variations in image quality can lead to discrepancies in the results. We would like to emphasize that this is a basic version of the application. Future improvements incorporating advanced machine learning algorithms and neural network techniques could further enhance application predictive capability. This could be a key focus in undertaking our future work.

| **TRIAL 1** | | **TRIAL 2** | |
|---|---|---|---|
| Blank Device Image | Test Device Image | Blank Device Image | Test Device Image |
| 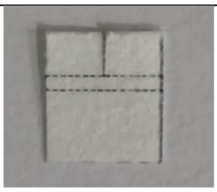 | 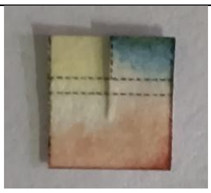 | 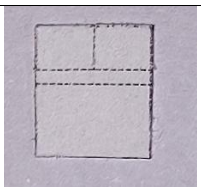 | 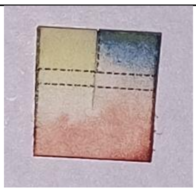 |

**References:**

Ardakani, F., Hemmateenejad, B., 2023. Pronounced effect of lamination on plasma separation from whole blood by microfluidic paper-based analytical devices. Anal. Chim. Acta 1279, 341767. https://doi.org/10.1016/j.aca.2023.341767

Kim, D., Kim, Sejin, Kim, Sanghyo, 2020. An innovative blood plasma separation method for a paper-based analytical device using chitosan functionalization. Analyst 145, 5491–5499. https://doi.org/10.1039/d0an00500b

Mukhopadhyay, M., Subramanian, S.G., Durga, K.V., Sarkar, D., DasGupta, S., 2022. Laser printing based colorimetric paper sensors for glucose and ketone detection: Design, fabrication, and theoretical analysis. Sensors Actuators B Chem. 371, 132599. https://doi.org/10.1016/j.snb.2022.132599

Nilghaz, A., Shen, W., 2015. Low-cost blood plasma separation method using salt functionalized paper. RSC Adv. 5, 53172–53179. https://doi.org/10.1039/c5ra01468a

Pokhrel, P., Jha, S., Giri, B., 2020. Selection of appropriate protein assay method for a paper microfluidics platform. Pract. Lab. Med. 21, e00166.

https://doi.org/10.1016/j.plabm.2020.e00166

Songjaroen, T., Dungchai, W., Chailapakul, O., Henry, C.S., Laiwattanapaisal, W., 2012. Blood separation on microfluidic paper-based analytical devices. Lab Chip 12, 3392–3398. https://doi.org/10.1039/c2lc21299d

Yang, X., Forouzan, O., Brown, T.P., Shevkoplyas, S.S., 2012. Integrated separation of blood plasma from whole blood for microfluidic paper-based analytical devices. Lab Chip 12, 274–280. https://doi.org/10.1039/c1lc20803a
